# Neurodynamic explanation of inter-individual and inter-trial variability in cross-modal perception

**DOI:** 10.1101/286609

**Authors:** G. Vinodh Kumar, Shrey Dutta, Siddharth Talwar, Dipanjan Roy, Arpan Banerjee

**Author notes:** Authors have equal contribution (G. Vinodh Kumar, Shrey Dutta). Joint corresponding authors (Dr Dipanjan Roy); (Dr Arpan Banerjee).

## Abstract

A widely used experimental design in multisensory integration is the McGurk paradigm that entail illusory (cross-modal) perception of speech sounds when presented with incongruent audio-visual (AV) stimuli. However, the distribution of responses across trials and individuals is heterogeneous and not necessarily everyone in a given group of individuals perceives the effect. Nonetheless, existing studies in the field primarily focus on addressing the correlation between subjective behavior and cortical activations to reveal the neuronal mechanisms underlying the perception of McGurk effect, typically in the “frequent perceivers”. Additionally, a solely neuroimaging approach does not provide mechanistic explanation for the observed inter-trial or inter-individual heterogeneity. In the current study we employ high density electroencephalogram (EEG) recordings in a group of 25 human subjects that allow us to distinguish “frequent perceivers” from “rare perceivers” using behavioral responses as well as from the perspective of large-scale brain functional connectivity (FC). Using global coherence as a measure of large-scale FC, we find that alpha band coherence, a distinctive feature in frequent perceivers is absent in the rare perceivers. Secondly, a decrease in alpha band coherence and increase in gamma band coherence occur during illusory perception trials in both frequent and rare perceivers. Source analysis followed up with source time series reconstructions reveals a large scale network of brain areas involving frontal, temporal and parietal areas that are involved in network level processing of cross-modal perception. Finally, we demonstrate that how a biophysically realistic computational model representing the interaction among key neuronal systems (visual, auditory and multisensory cortical regions) can explain the empirical observations. Each system involves a group of excitatory and inhibitory Hindmarsh Rose neurons that are coupled amongst each other. Large-scale FC between areas is conceptualized using coupling functions and the identity of a specific system, e.g., visual/ auditory/ multisensory is chosen using empirical estimates of the time-scale of information processing in these systems. The model predicts that the disappearance of alpha band coherence observed in rare perceivers stems from a negligible direct A-V (audio-visual) coupling however, an increase in indirect interaction via multisensory node leads to enhanced gamma band and reduced alpha band coherences observed during illusory perception. Overall, we establish the mechanistic basis of large-scale FC patterns underlying cross-modal perception.

## Introduction

Speech perception during face-to-face conversation inextricably involves multisensory integration of auditory and visual cues. This is nicely demonstrated in laboratory settings by the McGurk effect (McGurk & Macdonald, 1976), in which the video stimulus of a human speaker with the sound of /*ba*/ superimposed on the lip movements /*ga*/ is perceived by the listener as a completely different syllable /*da*/ (illusory/ cross-modal percept). Subsequently several studies have identified the psychophysical parameters that play a dominant role in eliciting cross-modal effects (Munhall et. al., 1996; van Wassenhove et. al., 2007, Thakur et al 2016) and their underlying neural mechanisms (Jones & Callan, 2003; Kaiser, 2004; van Wassenhove et. al., 2005; Saint-Amour et. al., 2007; Beauchamp, 2010; Keil et. al., 2012; Kumar et al 2017). Nonetheless, the distribution of responses to McGurk stimulus is heterogeneous and some individuals rarely perceive the illusion (Nath & Beauchamp, 2012a). While the neural correlates underlying illusory/ cross-modal perception has been extensively studied in a group of McGurk perceivers, the neurophysiology subserving the perceptual heterogeneity as well as the brain network mechanisms across individuals remains unclear.

Recent evidences show that subject-wise variability in the illusory perception is contingent on the McGurk stimulus and the response choice employed in the experimental paradigm (Mallick et. al., 2015). Concurrently, neuroimaging evidences attribute the heterogeneity across individuals to the extent of activation at the superior temporal sulcus (STS) (Beauchamp, 2010; Nath & Beauchamp, 2012b). Neurophysiological studies highlight the pre-stimulus activity in STS and its functional connectedness to front-parietal regions as a neuromarker of illusory perception within a group of individuals (Keil et al., 2012). More recent studies have indicated that beyond a specific region of interest, a large-scale network of oscillatory brain networks are involved in effectuating cross-modal perception(Kumar et al., 2016). A key question emerges how robust is this network across a group of individuals and whether the organization of these networks contingent on the stimulus configurations or the perceptual outcome, specifically in the case of McGurk incongruent stimulus. Secondly, what are the neural mechanisms that give rise to the network level correlates? While the first question needs to be answered empirically using a detailed neurophysiological study of underlying brain networks, the more broader question of systems-level understanding or functional brain network organization require a neurobiologically inspired computational model. The existing models of multisensory integration are either motivated from the context of response choices and probabilistic distribution of stimulus cues in the environment (Körding et al., 2007) or explanation of behavior from neurally inspired models (Thakur et al., 2016; Cuppini et al., 2017). Typically these models attempt to explain the firing rate dynamics of single neurons or the local population using a combination of synaptic and stimuli inspired parameters. Thus, the explanation of neurophysiological findings observed at the macroscopic scale of EEG and MEG remains elusive because of the dearth of a network model that captures the large-scale network dynamics.

In the current study we use the psychophysical variable of audio-visual (AV) lag that can modulate the degree of illusory perceptual experience in a group of individuals. We estimate the large-scale network underlying illusory perceptual experience in a group of individuals who frequently perceive McGurk illusion as well as investigate the functional network reorganization in individuals who rarely perceive the McGurk illusion. We find two distinct large-scale mechanisms operation during the multisensory information processing: 1) increase in gamma band global coherence and decrease in alpha band global coherence during illusory perception trials in both frequent and rare perceivers and 2) absence of peak in alpha band coherence across both illusory and unimodal perception trials in rare perceivers. Both these mechanisms were validated at the sensor level data and from source connectivity analysis using the LCMV beamformer (Van Veen et al., 1997). Subsequently, we designed a neural mass model that captures the global coherence dynamics observed in the EEG data. Previous studies have argued this kind of modeling is ideally suited to explain the emergence of spontaneous rhythmic patterns in EEG (Becker et al., 2015). Here we illustrate that a largescale model of multisensory interactions involving distinct local neuronal populations e.g., unisensory areas (Heschls’s gyrus/ STG and higher visual areas) and multisensory convergence zones (STS) can generate the synchronization patterns in sensor and source dynamics. Each local population consists of excitatory and inhibitory neural populations that are interconnected using biophysically observed parameters and each neuron within the population are capable of generating periodic spiking and bursting dynamics. Finally, we could illustrate how direct auditory-visual coupling whose presence was reported in neuroanatomical studies (Falchier et al., 2002; Rockland & Ojima, 2003; Wallace et al., 2004) and indirect interactions between audio-visual areas via multisensory convergence sites (Bizley & King, 2012) can bring forth distinct network mechanisms to facilitate perceptual experience.

## Results

### Inter-subject variability: Frequent and rare perceivers of illusory McGurk perception

We employed the incongruent McGurk stimulus, visual */ka/ paired with* auditory */pa/* to induce the illusory response /*ta*/. Overall, we used four kinds of AV stimuli: three McGurk incongruent pair with AV lags −450 ms (audio leads the articulation), 0 ms (synchronous), +450 ms (articulation leads the audio) and one congruent AV stimulus (visual */ta/* with auditory */ta/*). Following a forced choice paradigm, the participants reported if they heard /*ta*/, /*pa*/ or something else (others). Concurrently, the participants’ eye gaze behavior was recorded by an infra-red based eye tracking device. We characterized a participant as a ‘frequent perceiver’ (N=15) if they responded with 60% of */ta/* response to the McGurk incongruent stimulus at any lag, −450, 0 or +450 ms, failing which the participants were categorized as a ‘rare perceiver’ (N=10). **Figure 1B, C** illustrates the distribution of perceptual categorization responses in frequent and rare perceivers to the McGurk incongruent stimuli. At all AV lags, 80% of the rare perceivers reported /*ta*/ in <45% trials (see **Figure 1 - figure supplement 1**). We ran a repeated-measures two-way ANOVA on the percentage responses with AV lags and perceptual categories (*/ta/* and */pa/*) as the variables within each group of participants and use *p*<0.05 to evaluate statistical significance. For frequent perceivers, we observed that AV lags had no influence on the percentage responses, *F* (2, 89) = 0.84, *p* = 0.44. However, we observed a significant variation of percentage responses between the two perceptual categories, *F* (1, 89) = 19.90, *p* < 0.0001. Also, the interaction between perceptual categories and AV lags was significant, *F* (2, 89) = 29.83, *p* < 0.0001. For rare perceivers, no influence of AV lags was observed, *F* (2, 59) = 0.27, *p* = 0.76. However, variation of percentage responses between the two perceptual categories was significant, *F* (1, 59) = 64.47, *p* < 0.0001. Also, no significant interaction was observed between the perceptual categories and AV lags, *F* (2, 59) = 0.47, *p* = 0.66. We also performed paired Student’s t-test on the percentage of responses (/*ta*/ and /*pa*/) at each AV lag for frequent and rare perceivers and use statistical threshold of *p*=0.05 to evaluate significance. In frequent perceivers, we find significantly higher percentage of /*ta*/ responses at 0 ms (*t* (14) = 7.81, *p* < 0.0001) and +450 ms AV lag (*t* (14) = 2.12, *p* = 0.04). No significant difference was observed at −450 ms (*t* (14) = 1.97, *p* = 0.06) AV lag. However, in rare perceivers we observed a significantly higher percentage of /*pa*/ responses at −450 ms (*t* (9) = −3.62, *p* = 0.002), 0 ms (*t* (9) = −4.93, *p* < 0.0001) and +450 ms (*t* (9) = −5.61, *p* < 0.0001) AV lag. Unpaired Student’s t test employed to compare the percentage of /*ta*/ responses during the congruent /*ta*/ stimulus showed no significant difference between the groups (*t* (23) = 2.02, *p* = 0.05) **Figure 1 - figure supplement 2**.

**Figure 1:**
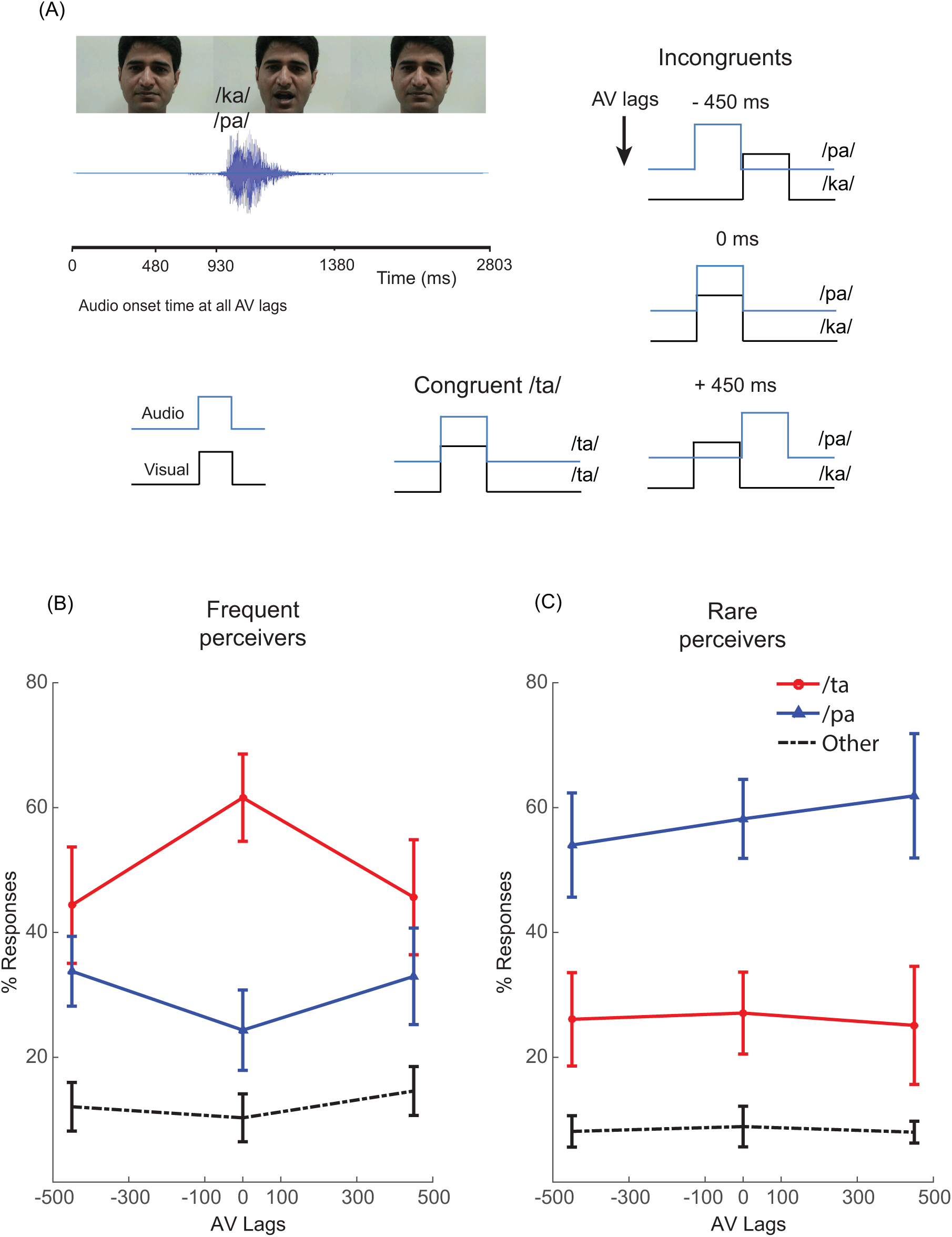
Experimental setup and behavior: (A) Video frames from the stimulus showing neutral face at the stimulus onset and the facial gesture during articulation (B) The McGurk stimuli: Audio /pa/ superimposed onto the lip movement /ka/ presented with AV lags −450 ms, 0 ms and +450 ms and the congruent stimulus: Audio /ta/ superimposed onto the lip movement /ta/. The location of the onset of the audio is place with respect to the articulator’s initiation of lip movement. (C) Group percentage distribution of the perceptual responses (/ta/, /pa/ and others) in frequent and rare perceivers.

Gaze fixations on the head and mouth areas of the speaker in the AV stimuli were converted into percentage measures for each subject on a trial-by trial basis and sorted based on the stimulus type and perceptual categories. The bar graphs in **Figure 1-figure supplement 3** illustrates the mean and the standard error of the percentage of gaze fixations on the mouth of the articulator during /*ta*/ and /*pa*/ perception averaged across the participants. We performed a repeated-measures two-way ANOVA on the percentage responses with AV lags and perceptual categories (*/ta/* and */pa/*) as the variables in frequent and rare perceivers. In frequent perceivers **Figure 1 - figure supplement 3A**, we observed that there was no influence of AV lags, *F* (2, 89) = 0.36, *p* = 0.70 and perceptual categories, *F* (2, 89) = 3.88, *p* = 0.05 on the percentage of gaze fixations on the mouth. Furthermore, the interaction effect between them was also insignificant, *F* (2, 89) = 0.07, *p* = 0.93. Similarly, in rare perceivers **Figure 1-figure supplement 3B**, AV lags, *F* (2, 59) = 2.54, *p* = 0.09 and perceptual categories, *F* (2, 59) = 0, *p* = 0.97 had no effect on the percentage of gaze fixations at the mouth. Also, no evidence of an interaction effect between them was observed, *F* (2, 59) = 0.2, *p* = 0.82. We further performed unpaired Student’s t-test to compare the percentage of gaze fixations on mouth between frequent and rare perceivers i.e frequent /*ta*/ vs Rare /*ta*/ and frequent /*pa*/ vs Rare /*pa*/. We observed that frequent perceivers elicited significantly higher percentage of fixations at mouth during /*ta*/ perception at −450 ms (*t* (23) = 3.42, *p* = 0.002), 0 ms (*t* (23) = 3.88, *p* = 0.0007) and +450 ms (*t* (22) = 2.79, *p* = 0.01) AV lag. Similarly, during /*pa*/ perception frequent perceivers elicited higher percentage of fixations on mouth at −450 ms (*t* (23) = 4.56, *p* < 0.001), 0 ms (*t* (23) = 2.95, *p* = 0.0071) and +450 ms (*t* (23) = 2.45, *p* = 0.02) AV lag.

### Large-scale functional connectivity dynamics

To investigate the underlying differences in dynamic functional connectivity (FC) between the perceptual categories we computed the global coherogram during /*ta*/ and /*pa*/ perception. Global coherogram defined from the normalized vector sum of all pairwise coherences amongst EEG sensors captures the evolution of global coherence in time and frequency domain simultaneously (Lachaux et al., 1999). Mathematically, global coherence is the ratio of the largest eigenvalue of the cross-spectral matrix to the sum of its eigenvalues (Mitra & Bokil, 2008). Subsequently, we compared the global coherogram of */ta/* and */pa/* at AV lags: −450ms (**Figure 2A, C**), 0ms (**Figure 2E, G**) and +450 (**Figure 2I, K**) using cluster based permutation tests. The onset of first stimulus was considered the point of reference for time-locking (zero). Positive clusters highlighted in black dashed rectangles and negative clusters in red dashed boxes signify time-frequency islands of increased and decreased synchrony respectively in the large-scale functional network. We also compared the presence of band-specific peaks/enhancement in global coherence in the frequent (**Figure 2B, F, J)** and rare perceivers (**Figure 2D, H, L)** during /*ta*/ and /*pa*/ perception using Silverman’s bootstrapping test for examining multimodality.

**Figure 2:**
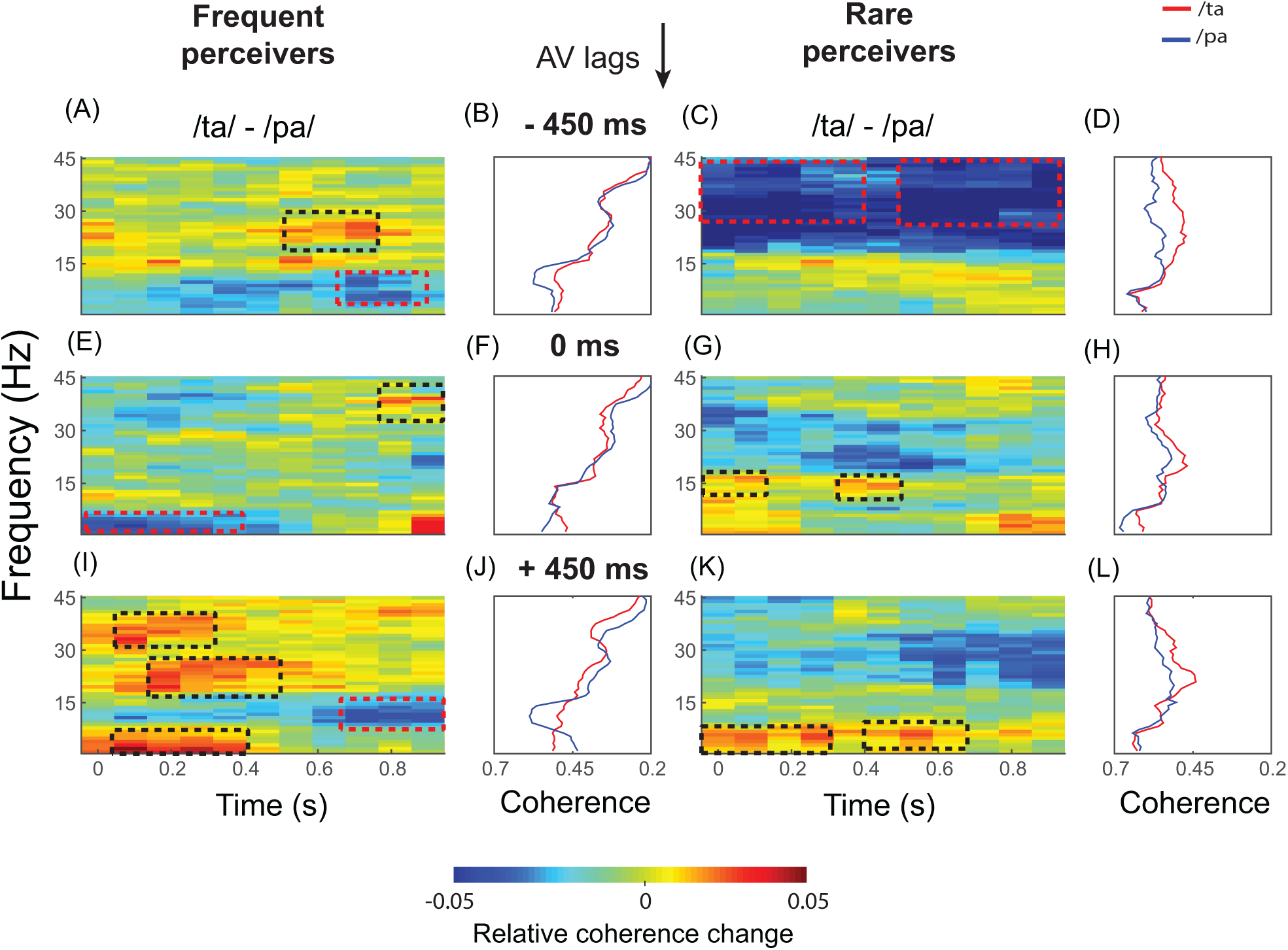
Large scale functional connectivity dynamics observed in sensor time series: Global coherogram differences between the perceptual categories (/ta/ and /pa/) and time averaged global coherence respectively during /ta/ and /pa/ perception in frequent and rare perceivers at −450 ms (A,B,C,D), 0 ms (E,F,G,H) and +450 ms (I,J,K,L) AV lag.

For frequent perceivers, during −450 ms AV lag (**Figure 2A**), we observed a negative cluster in the theta (*z*_0.05_ = −5.31) in the temporal range of 650-800 ms and a positive cluster in the beta band (*z*_0.95_ = −3.84) between 500 ms to 700ms. For videos at 0 ms AV lag (**Figure 2E**), we observed a negative cluster in the theta (*z*_0.05_ = −6.05) and alpha band (*z*_0.05_ = −5.81) in the temporal range of ∼0-450 ms, and a positive cluster (*z*_0.95_ = −4.32) in gamma band between 800 and 900 ms. During +450 ms AV lag (**Figure 2I**), we observed three positive clusters, (1) in the theta band (*z*_0.95_ = −5.52) in the 100-400 ms time window, (2) in the beta band (*z*_0.95_ = −4.40) between 200 to 500 ms, and (3) in the gamma range (*z*_0.95_ = −4.34) in the temporal window of 50 ms and 250 ms. Also, a negative cluster in the alpha band (*z*_0.05_ = −5.85) was observed between 700 to 900 ms time window.

For rare perceivers, at −450 ms AV lag (**Figure 2C**), we observed two prominent negative clusters (*z*_0.05_ = −6.72) spanning gamma band in the temporal range of 0-400 ms and ∼500-900 ms. For videos with 0 ms AV lag (**Figure 2G**), we observed two positive clusters in the beta band (*z*_0.05_ = −5.69) and (*z*_0.05_ = −5.68) between ∼0-150ms and ∼300-500 ms respectively. At +450 ms AV lag (**Figure 2K**), we observed two positive clusters (*z*_0.95_ = −6.16) and (*z*_0.95_ = −6.04) in the theta band in the temporal window of ∼0-300 ms and ∼400-700 ms respectively.

Cluster based permutation tests, performed to test the differences in global coherogram between /*ta*/ and /*pa*/ elucidated the neural signatures in large-scale FC corresponding to inter-trial variability observed within frequent and rare perceivers. Consequently, to address if inter-individual heterogeneity stems from the differences in the inherent processing of multisensory stimuli in the two groups of perceivers, we evaluated if any frequency specific enhancement of global coherence occurs during /*ta*/ and /*pa*/ perception. Consequently, we computed the global coherence during /*ta*/ and /*pa*/ perception for frequent and rare perceivers that would provide a holistic picture of the underlying large-scale FC. In frequent perceivers we observed qualitatively that the global coherence followed a bimodal distribution during /*ta*/ and /*pa*/ perceptions across all AV lags (**Figure 2B, F, J**). The modes were primarily centered around alpha (8-13 Hz) and gamma (30-40 Hz) bands, signifying enhanced coherence. Silvermann’s bootstrapping test employed to examine the statistical significance of those peaks revealed significant bimodal peaks (*p* < 0.05) during /*ta*/ and /*pa*/ perception at −450 ms, 0 ms AV lags. However, there were no significant bimodal peaks during /*ta*/ *(p* = 0.07) and /*pa*/ perception *(p* = 0.13) at +450 ms AV lag (**Figure 2J**).

In rare perceivers, Silvermann’s bootstrapping test revealed significant bimodal peaks only during /*ta*/ perception at 0 ms AV lag (*p* < 0.05) (**Figure 2H**). There were no significant bimodal peaks during /*pa*/ perception at −450 ms (*p* = 0.35) (**Figure 2D**), 0 ms (*p* = 0.30) (**Figure 2H**) and +450 AV lag (*p* = 0.15) (**Figure 2L**). Similarly, no significant bimodal distribution were observed during /*ta*/ perception at −450 ms (*p* = 0.21) (**Figure 2D**) and +450 ms (*p* = 0.20) AV lags (**Figure 2L**). Importantly, the bimodal peaks during /*ta*/ perception at 0 ms AV lag were clustered around delta (1-4 Hz), theta (4-8 Hz) and gamma (30-40 Hz). Notably, a desynchronization in the alpha band was observed in rare perceivers (**Figure 2D, H, L**) across all AV lags and perceptual categories.

To further understand if these frequency specific coherence differences contingent on the stimulus configurations, we computed the global coherogram and time averaged global coherence during congruent /*ta*/ in frequent and rare perceivers and compared them using cluster based permutation tests (**Figure 3**). Global coherogram differences in congruent /*ta*/ between frequent and rare perceivers computed employing cluster based permutation tests revealed three negative clusters, (1) in the beta band between ∼50-400 ms (*z*_0.05_ = −5.87) temporal window, (2) in the beta band between the time window ∼700-900 ms (*z*_0.05_ = −5.79) and (3) in the gamma band from ∼150-900 ms (*z*_0.05_ = −6.54). A positive cluster in the delta and theta band was also observed between ∼700-900 ms (*z*_0.95_ = −6.24) time window (**Figure 3C**). Conspicuously, the global coherence distribution for the congruent /*ta*/ in frequent and rare perceivers followed a similar pattern (**Figure 3D**).

**Figure 3:**
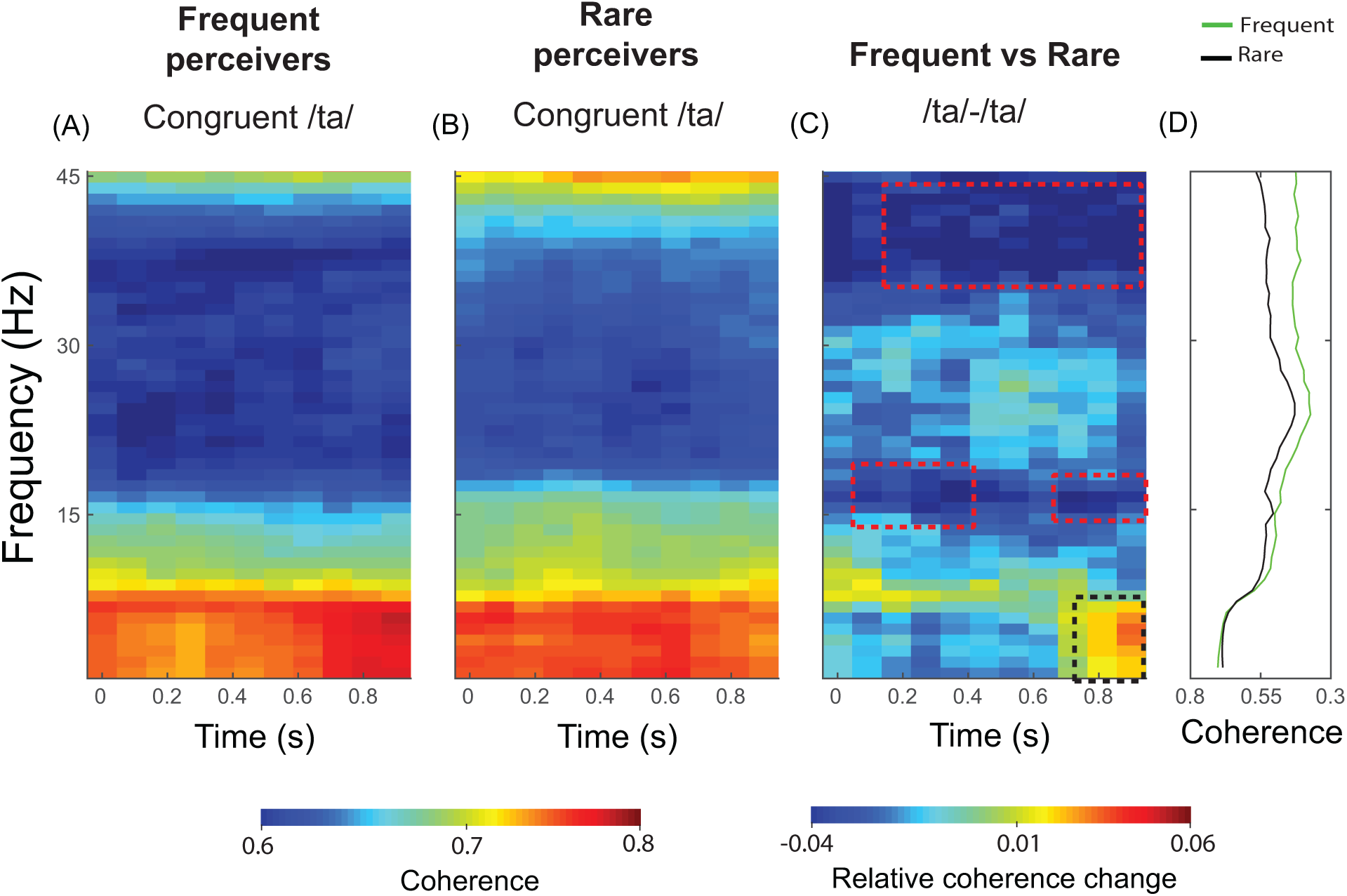
Large-scale functional connectivity dynamics during congruent /ta/: (A) Global coherogram during congruent /ta/ perception in (A) Frequent perceivers (B) Rare perceivers (B) Global coherogram difference between frequent and rare perceivers (D) Time averaged global coherence during /ta/ and /pa/ perception in frequent and rare perceivers.

### Source analysis reveals cortical areas participating in functional connectivity dynamics

To validate the role of the identified sources in the overall functional connectivity pattern observed in the sensor EEG, we initially identified the cortical generators of the EEG time series by employing linear constrained minimum variance (LCMV) beamformer algorithm (Van Veen et al., 1997). Subsequently, we projected the epoched time series into the source time space by multiplying them with the concordant spatial filter (constructed by LCMV beamformer, for more info. see methods) of the source locations that showed statistical significance in the ratio of source power between /*ta*/ and /*pa*/ trials. Finally, we computed the global coherogram for the perceptual categories and compared them using cluster based permutation tests. Elicitation of a similar trend in the global coherogram differences essentially confirms the involvement of the identified sources in the large-scale FC underlying McGurk perception. The sources eliciting statistical significance in the ratio of source power between /*ta*/ and /*pa*/ are illustrated in **Figure 4A** and the source locations are listed in **Table 1**. The source locations were consistent across all AV lags and between frequent and rare perceivers. Cluster based permutation tests employed to compare the global coherogram (/*ta*/ -/*pa*/) computed from the source time series revealed in frequent perceivers at −450 ms AV lag (**Figure 4B**) one positive cluster in the alpha band (*z*_0.95_ = −5.18) in the temporal range of ∼200-700 ms,. During 0 ms AV lag (**Figure 4D**), three negative clusters, two clusters in the alpha band in the temporal window of ∼0–100ms (*z*_0.05_ = −6.03), ∼600-900ms (*z*_0.05_ = −6.05) and one (*z*_0.05_ = −6.93) in the low gamma band between ∼150-350 ms. At +450 ms AV lag (**Figure 4F**), one prominent positive cluster in the high beta and gamma band (*z*_0.95_ = −6.65) spanning the entire stimulus duration, and two negative clusters spanning the theta and alpha band in the time window of ∼0-400 ms (*z*_0.05_ = −5.57) and between ∼500-650 ms (*z*_0.05_ = −5.62) was observed.

**Table 1:** The table lists the cortical loci that elicited power higher than the set threshold (> 99.5 percentile) in the source analysis

|  | Left hemisphere | Right hemisphere |
| --- | --- | --- |
| **Frontal lobe** | Inferior frontal gyrus<br>Middle frontal gyrus<br>Superior frontal gyrus<br>Cingulate gyrus | Inferior frontal gyrus<br>Middle frontal gyrus<br>Superior frontal gyrus<br>Cingulate gyrus |
| **Temporal lobe** | Fusiform gyrus<br>Middle temporal gyrus<br>Superior temporal gyrus | Fusiform gyrus<br>Middle temporal gyrus<br>Superior temporal gyrus |
| **Parietal Lobe** | Precuneus |  |

**Figure 4:**
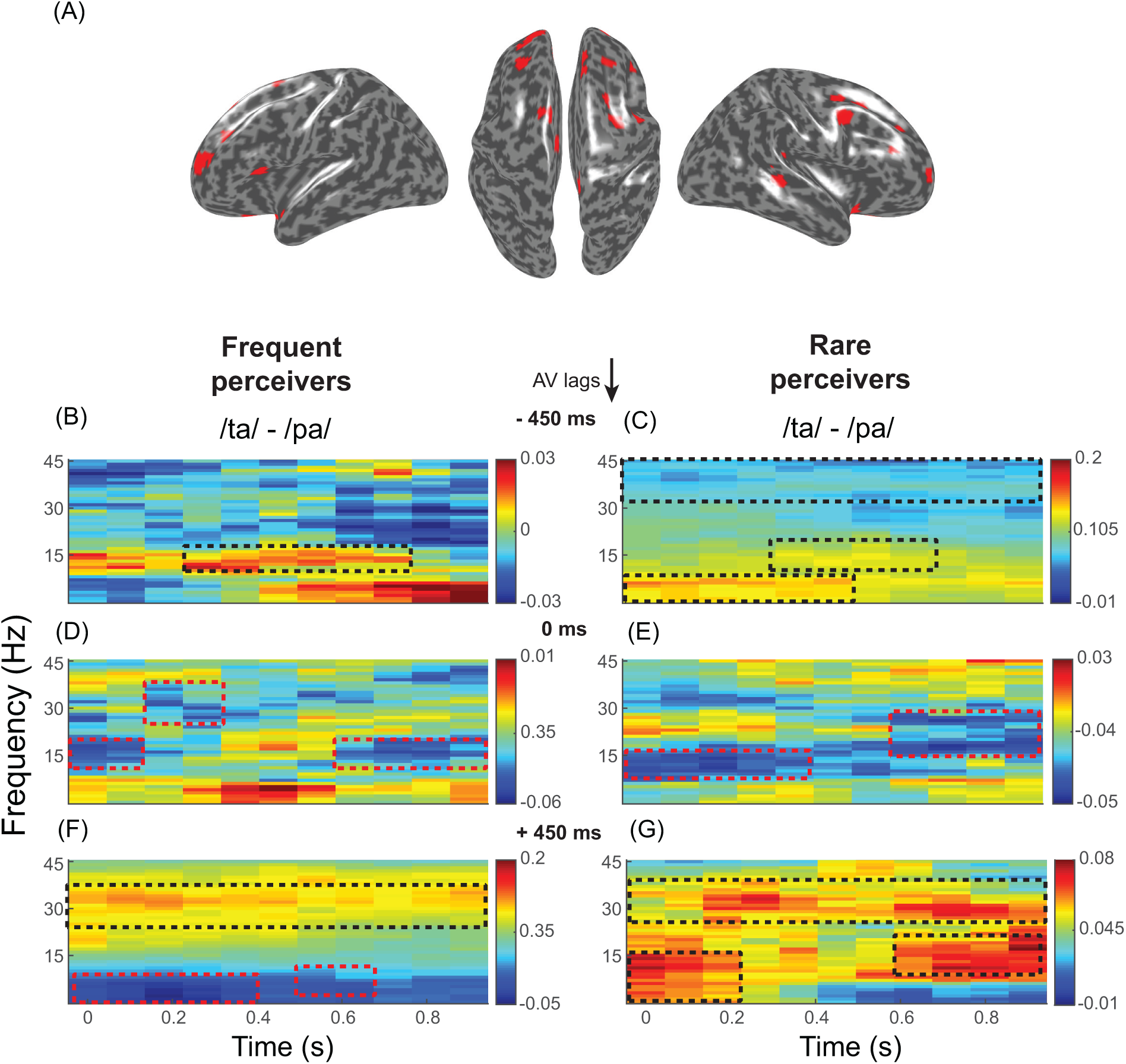
Source reconstruction: (A) Sources identified using the LCMV beamformer algorithm from the sensor time series. The source power of the ratio between /ta/ and /pa/ eliciting power than the set threshold (¿99.5 percentile) are highlighted. Global coherogram differences between the perceptual categories (/ta/ and /pa/) computed from the source-time series in frequent and rare perceivers during −450 ms (B,C), 0 ms (D,E) and +450 ms (F,G) AV lag.

For rare perceivers, during −450 ms AV lag (**Figure 4C**), three positive clusters, (1) in the theta and alpha band (*z*_0.95_ = −4.56) from the onset to ∼500ms, (2) in the beta band between ∼300-700 ms and (3) a prominent positive cluster (*z*_0.95_ = −4.56) in the gamma band spanning the entire stimulus duration was observed. At 0 ms AV lag (**Figure 4E**), a negative cluster (*z*_0.05_ = −5.39) in the alpha band between from stimulus onset to ∼600 ms and a negative cluster (*z*_0.95_ = −4.57) in the beta band in the temporal range of ∼600-900 ms was observed. During +450 ms AV lag (**Figure 4G**), three positive clusters, (1) in the theta band and alpha band (*z*_0.95_ = −4.60) between ∼0-220 ms, (2) in the beta band (*z*_0.95_ = −4.16) between ∼600-900 ms, and (3) in the gamma band (*z*_0.95_ = −4.38) spanning the entire stimulus duration was observed.

### Network model comprising of 3 neural masses with fast, intermediate and slow time-constants generates alpha and gamma coherence

We incorporated a neural mass model approach (Becker et al., 2015; Aerts et al., 2018) to investigate the alpha and gamma coherence dynamics associated with inter-individual and inter-trial variability respectively. Since EEG data does not necessarily reflect the local synaptic activity, neural mass model which operates to phenemenologically explain mesoscopic and macroscopic features in EEG/ MEG data offers an attractive tool to understand the underlying neural mechanisms (Lopes da Silva et al., 1974; Jansen & Rit, 1995; David & Friston, 2003). A neural mass is essentially an abstraction of summed synapto-dendritic activity of several thousand neurons in an area which can be in a cooperative dynamical state such as synchronous firing that gives rise to low-frequency oscillations. Such shared dynamical states allow us to reduce the population dynamics in terms of coupled ordinary differential equations where explicit spatial effects can be ignored (Stefanescu & Jirsa, 2008). Armed with the knowledge of cortical sources underlying cross-modal perception (**Table 1**) we consider broadly a network of three neural masses as the underlying neuro-cognitive network comprising of auditory, visual and cross-modal masses (nodes). Each node can be further expanded as a population of excitatory and inhibitory Hindmarsh-Rose (HR) neurons (Hindmarsh & Rose, 1984) representing auditory, visual and multisensory areas. The key parameters that govern the time scale of the oscillatory dynamics come from physiologically motivated parameter values for each neural area. For instance, the auditory node is assumed to be the most sensitive to ambient temporal fluctuations hence operating with a fast time-scale, visual node the slowest in terms of sensitivity and somewhat intermediate time-scale for multisensory node (see materials and methods for details, **Figure 5**). The existence of two time-scales facilitates the co-existence of synchronous states in alpha and gamma oscillations when slow (visual) node is source of excitatory influence (EI) and fast (auditory) node is sink of EI and when coherence was computed across all nodes. These co-existent states emerge via two possible routes, 1) when visual node (V) interacts with the auditory node (A) through direct coupling (*W*_*AV*_) and 2) when indirect coupling (*W*_*AM*_ &*W*_*VM*_) between A-V nodes via the multisensory node (M) range from 0.35 to 0.7 (**Figure 6 Supplement 1A**). We assume coupling strength less than 0.35 to be weak coupling (WC), coupling strength between 0.35 and 0.7 to be moderate coupling (MC) and coupling greater than 0.7 to be strong coupling (SC). We also observe high coherence around alpha band and gamma band in SC range however, a distinct peak around alpha band is not clearly observed. Any other model configuration is not able to create the co-existence of alpha and gamma band coherence in MC range (**Figure 6 Supplement 2**). Further, when the fast-slow interaction takes place via direct coupling alone (*W*_*AV*_ ranges from 0 to 1, *W*_*AM*_ &*W*_*VM*_ = 0) we observe the existence of only alpha band coherence but not the gamma band coherence (**Figure 6 Supplement 1B**). Here, the absence of gamma band coherence implies a diminished indirect coupling of A-V nodes via multisensory node (*W*_*AM*_ &*W*_*VM*_). Moreover, we observe only gamma band coherence in MC range (**Figure 6 Supplement 1C**) when we restricted the fast-slow (A-V) interactions via multisensory node alone (indirect A-V coupling *W*_*AM*_ &*W*_*VM*_ range from 0 to 1, *W*_*AV*_ = 0). This observation clearly links alpha coherence to direct A-V coupling whereas gamma coherence to indirect A-V coupling (A-M-V) of neural masses.

**Figure 5:**
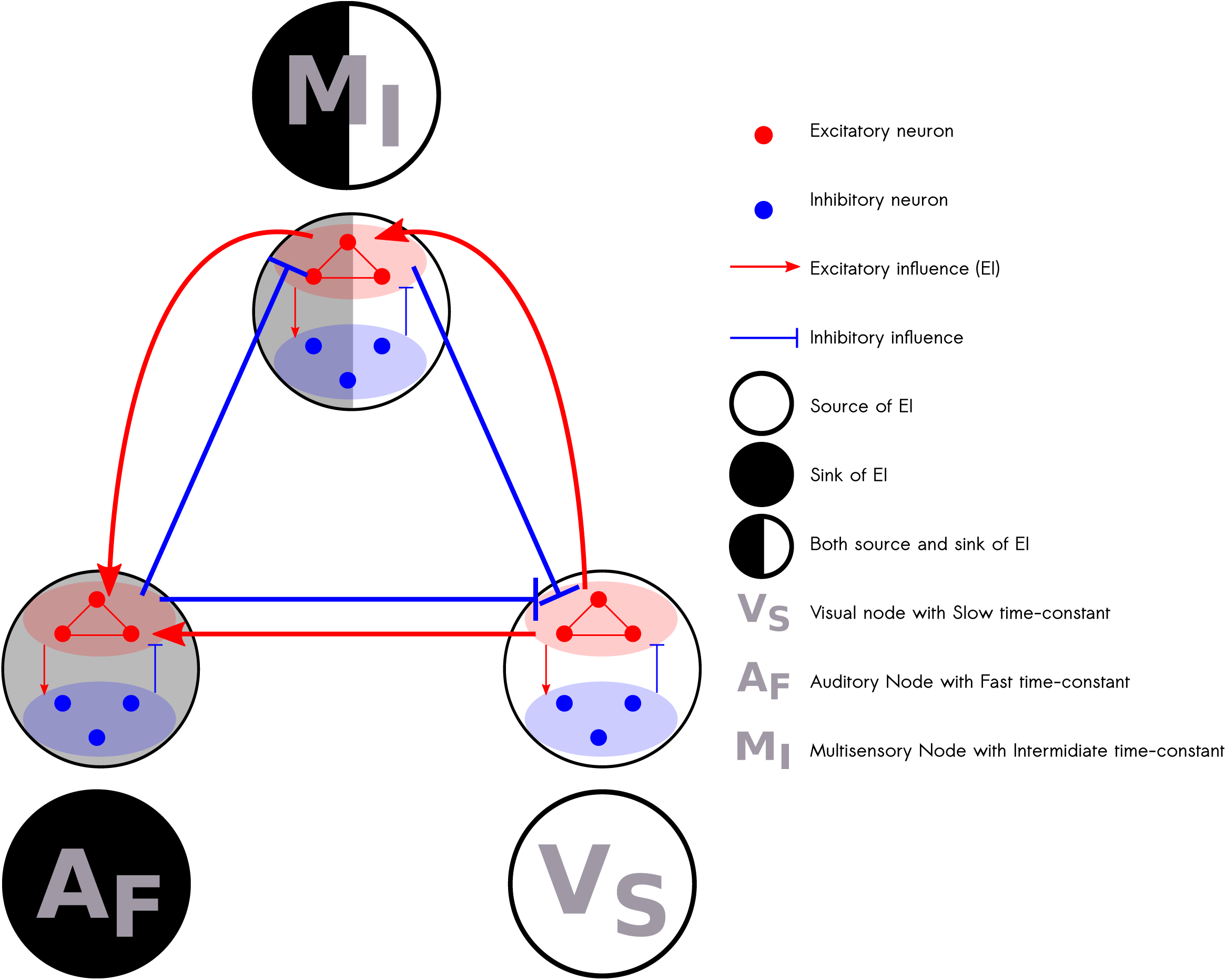
Large scale dynamical model consisting of a network three neural masses with different time-constants: The model comprises three nodes representing auditory (fast time-constant), visual (slow time-constant) and higher order multisensory regions (intermediate time-constant). Each node consists of network of 100 Hindmarsh-Rose excitatory and 50 inhibitory neurons. Each neuron can exhibit isolated spiking, periodic spiking and bursting behavior. Exci-tatory influences between the nodes are balanced by their inhibitory counterpart. The source and sink represent the flow of excitatory influence.

**Figure 6:**
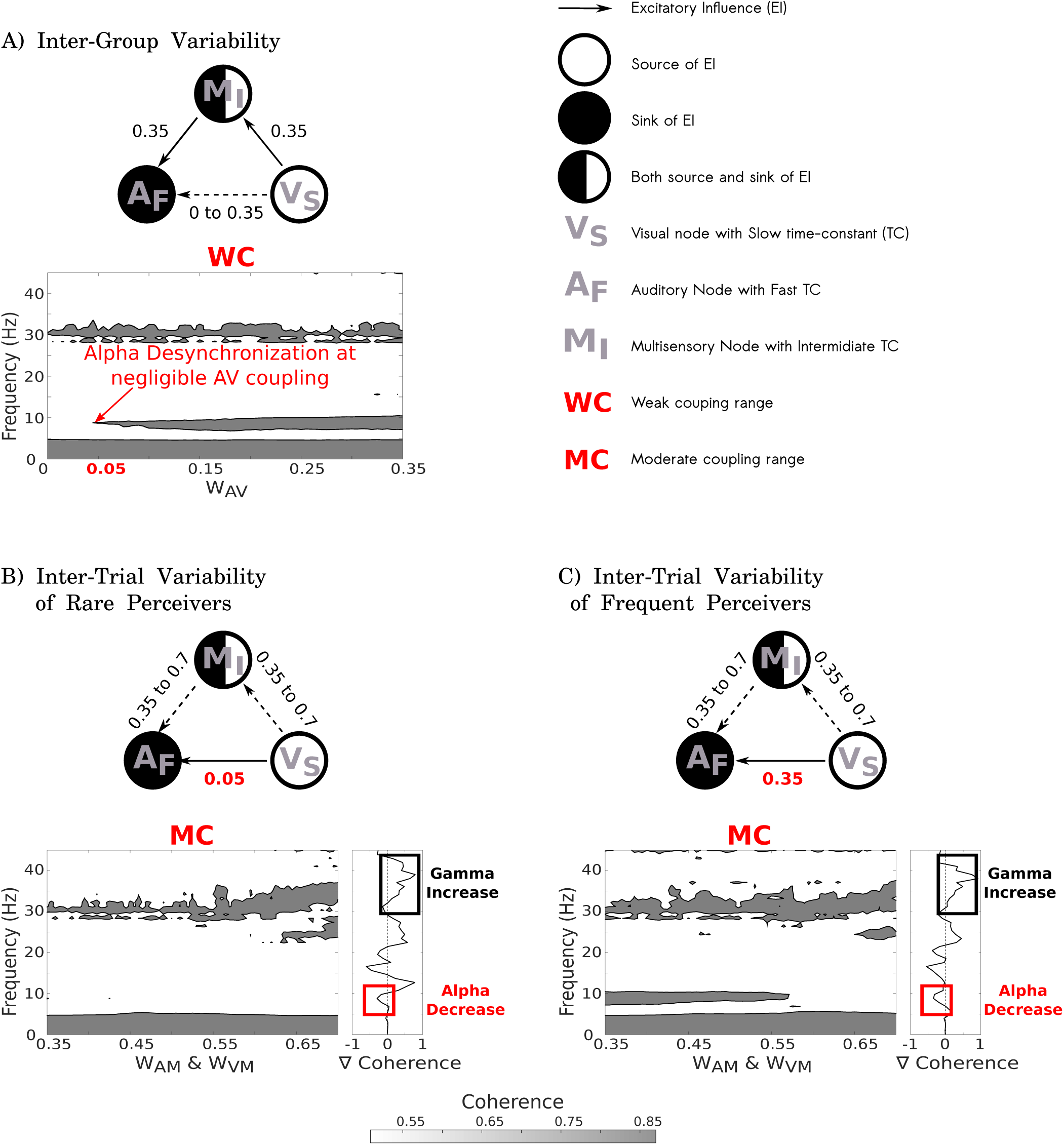
Mechanistic understanding of Inter-individual and inter-trial variability: A) Alpha de-synchronization characteristic of rare perceivers resulted due to negligible A-V coupling. B) & C) Enhanced gamma coherence and reduced alpha coherence observed in illusory perception is due to increase in indirect coupling involving multisensory node irrespective of the influence of direct A-V coupling.

### Direct Audio-Visual interaction underpins Inter-Individual Variability

Our empirical results suggest that negligible alpha coherence is a hallmark of rare perceivers. Since, direct A-V interaction generates a peak around alpha coherence (**Figure 6 Supplement 1 A & B**), we hypothesize that lesser amount of direct interaction or even absence of it is associated with de-synchronization of alpha band coherence. To test this hypothesis, we start with a balanced network coupling state, *W*_*AV*_ = *W*_*AM*_ = *W*_*VM*_ = 0.35, where alpha and gamma band coherences co-exist (**Figure 6 Supplement 1A**) and study the change in the coherence peaks as direct A-V coupling (*W*_*AV*_) decreases. In **Figure 6A**, we observe a suppression of alpha coherence peak as A-V coupling decreases; however gamma coherence peak remains more or less intact. Further, when A-V coupling becomes negligible (*W*_*AV*_ < 0.05) we observe disappearance of alpha coherence peak. This suggests that alpha de-synchronization can stem from low direct A-V coupling in rare perceivers.

### Audio-Visual interaction via Multisensory node underpins Inter-Trial Variability

Broadly speaking, enhanced gamma coherence and decreased alpha coherence is observed unequivocally in frequent perceivers and rare perceivers when illusory and non-illusory trial comparisons were extracted to study the inter-trial variability. Even though rare perceivers exhibited overall lower alpha coherence, the differential decrease in alpha band coherence was clearly observed at sensor and source level (**Figure 2 & 4**). As shown earlier, decrease in direct A-V coupling causes a decrease in alpha band coherence (**Figure 6A**) in rare perceivers and hence decrease in direct A-V coupling cannot be associated with illusory perception. However, gamma band coherence peaks emerge as a coexistent state once indirect A-V interactions via multisensory node are incorporated in the model (**Figure 6 Supplement 1 A & C**) allowing us to propose a dominant role of interactions between multisensory and unisensory areas modulating cross-modal perception. To test this hypothesis for frequent perceivers we start with a balanced network configuration that generates co-existing alpha band and gamma band coherence (*W*_*AV*_ = *W*_*AM*_ = *W*_*VM*_ = 0.35, **Figure 6 Supplement 1A**) and for rare perceivers we choose a network configuration that generates peak only around gamma band (*W*_*AV*_ = 0.05;*W*_*AM*_ = *W*_*VM*_ = 0.35, **Figure 6A** & **Figure 6 Supplement 1C**). Then, we track the change in gamma coherence as indirect A-V interaction via multisensory node (*W*_*AM*_ &*W*_*VM*_) increases simultaneously in MC range (0.35 to 0.7). As hypothesized, we observe an increase in gamma coherence in network configurations for both frequent and rare perceivers. Interestingly, increasing indirect A-V interactions not only increases gamma band coherence but also display a decrease around alpha band coherence in network configurations of frequent as well as rare perceivers even though rare perceivers exhibit overall weaker alpha band coherence (**Figure 6 B & C**). Thus, our model implicates an increase in indirect A-V interaction via multisensory node leading to an increase in gamma band coherence as well as a decrease in alpha band coherence and thus facilitating illusory perception.

## Discussion

A vast body of work has used the Mcgurk paradigm to study cross-modal perception and the numbers are only increasing (Alsius et al., 2018). An ongoing challenge still remaining to the community is accurate identification and characterization of possible neural mechanisms that govern the behavioral variability. For example, why do some people perceive it so strongly, whereas others do not? An approach taken by brain stimulation studies had earlier addressed the issue of inter-individual variability, and identified the candidate brain areas that are probably responsible (Beauchamp, 2010). A more emerging understanding suggest the existence of networks of brain regions facilitating perceptual processing (Bressler & Menon, 2010), nonetheless the neurophysiological correlates of inter-individual variability are yet to be understood. In this perspective, a recent review suggests neuronal oscillations as a key substrate of neuronal information processing that needs to be fully explored to answer the individual’s perceptual experience (Keil & Senkowski, 2018). It is well known that robust oscillations observed from macroscopic recordings such as EEG/ MEG are an outcome of network interactions among local subpopulations of excitatory and inhibitory neurons (Wilson & Cowan, 1972; Deco et al., 2010; Becker et al., 2015). Empirically such interactions result in global coherence dynamics observed by earlier studies such as Kumar et al (Kumar et al., 2017). In the current study we demonstrate how distinct coherence patterns further become the hallmark of category specific perceptual experience such as the presence of alpha band coherence became a group-labeling attribute for perceptual categorization. Furthermore we find that across trials, the pattern of coherence dynamics determine the trial-specific perceptual outcome. Finally, using computational models of interactive large-scale brain networks, we capture the neural mechanisms through which coherence dynamics evolve in the brain. Put together, we present an attractive mechanistic proposal that underlie the observed inter-individual and inter-trial variability in multisensory speech perception.

The key empirical observations in our study are: (1) Rare perceivers exhibit a diminished alpha band global coherence, indicating desynchronization of large-scale neural assemblies in the alpha band (2) Both rare and frequent perceivers’ cross-modal perception (such as */ta/)* involves an enhanced gamma band coherence and decrease in alpha band coherence compared to unimodal perception (such as */pa/)*. The large-scale neuro-dynamic model of cross-modal perception suggests de-synchrony in the alpha band, characteristic of rare perceivers, is due to extremely weak direct A-V coupling (*W*_*AV*_ < 0.05). Furthermore, an increase in indirect interaction between auditory and visual systems via multisensory node (increase in *W*_*AM*_ &*W*_*VM*_) facilitates high level of synchronization in gamma band and a desynchronization at alpha band. We further elaborate on our empirical and modeling results in the following subsections.

### Heterogeneous nature of illusory perception

Trial-by-trial variation of perceptual experience within an individual has been previously reported by several studies (Beauchamp, 2010; Keil et al., 2012; Roa Romero et. al., 2015, Kumar et. al. 2016). Behavioral results (**Figure 1B, C**) also indicate that the entire population of volunteers can be distinctly classified in two categorical groups: frequent perceivers and rare perceivers. Similar inter-individual variability were observed and quantified by previous studies (Nath & Beauchamp, 2012; Proverbio et al., 2016). We also presented the McGurk incongruent video (/*pa*/-/*ka*/) with varying temporal asynchrony, AV lags of ±450ms. Perceptual experience of frequent perceivers was modulated as a function of lags, however, no such effect was observed in rare perceivers. The decrease in McGurk perception for ±450ms AV lags is consistent with the existing studies (Munhall et al., 1996; van Wassenhove et al., 2007). Also for ±450ms AV lagged videos, higher degree of illusory perception was observed in frequent perceivers compared to rare perceivers. Furthermore, irrespective of the perception (/*ta*/ or /*pa*/) the gaze fixations on the mouth of the articulator were also significantly lower in rare than frequent perceivers. The distinctness in the behavior of rare perceivers pinpoints a difference in the processing of multisensory speech. Therefore, we expected to identify neurophysiological correlates that can characterize a rare perceiver from the frequent perceivers as well as the cross-modal perceptual experience from the unimodal perception that varies trial-by-trial within an individual. Ideally, a single measure that captures these different kinds of heterogeneity, inter-individual and inter-trial can set the ideal platform for discussing about network mechanisms.

### Neuromarkers of inter-individual and inter-trial variability

Large-scale systems of distributed and interconnected neuronal populations organized to perform specific cognitive tasks are referred to as neurocognitive networks (NCNs) (Bressler & Menon, 2010). Multisensory speech perception that requires the integration of information among spatially distinct sensory systems, components of which are often distributed over the whole brain becomes an ideal candidate to explore from the perspective of NCNs. In physiological signals NCNs can be studied by quantifying the extent of coordination among neuronal assemblies over the whole brain (Bressler, 1995; Bressler & Kelso 2001). The most significant achievement of our study was to capture the network correlates of inter-individual and inter-trial variability with the same measure of global coherence at both sensor level and source level EEG analysis. Our results show that frequent perceivers exhibit enhanced global coherence in the alpha band than rare perceivers. Notably, the enhancement was consistent across all AV lags in frequent perceivers. Previous evidences accentuate the modulations in alpha band coherence to central executive processes (Klimesch, 1999; Sauseng et. al., 2005) that are postulated to be involved in allocating working memory storage to phonological loop that maintains verbal information, and the visuo-spatial sketchpad that maintains transient visuo-spatial information (Baddeley, 1992). Therefore, we posit that the enhanced global coherence in alpha band as a marker that characterizes the presence of specific NCN level processing in frequent perceivers which is absent in rare perceivers.

Recent study by Fernández and colleagues demonstrates an increase in the power of theta oscillations in response to an incongruent McGurk stimulus accentuating its role in the prediction of the conflict (Fernández et al., 2018). Noticeably, we observed an enhanced global coherence in the theta band in frequent and rare perceivers irrespective of the perceptual experience which indicates even if theta band communication is present in both group of perceivers, it is a not necessarily a marker of inter-individual differences or trial specific perception. In general it is quite possible that different neuro-cognitive processes can be operating simultaneously involving communication at various frequencies via coherence (Senkowski et al., 2008). Hence, it is important to identify which of these are meaningful to the ongoing task and the subtle differences that vary with the context in which the task evolves. In an earlier study Kumar et al. (Kumar et al., 2016) have showed that global coherogram captures the difference in processing of crossmodal (illusory /*ta*/) and unimodal (non-illusory /*pa*/) perception in frequent perceivers from a subset of data that we present in this manuscript. While the detailed pattern of coherogram differences between /*ta*/ and /*pa*/ trials in perceivers and rare perceivers are slightly different, there was an enormous similarity in trend of coherence differences in distinct spectro-temporal locations that was conspicuous. For example, both frequent and rare perceivers have enhanced gamma band coherence and diminished alpha band coherence in /*ta*/ trials compared to /*pa*/ trials for temporally synchronous AV stimuli. For asynchronous trials, broadband coherence enhancement in both frequent and rare perceivers was observed. Based on these observations we argue that global coherogram differences (/*ta*/-/*pa*/) present itself as a signature of the inter-trial perceptual variability. Furthermore, frequency specific signature in the global coherence consistent across the perceptual categories enhanced alpha and gamma band coherence in frequent perceivers and desynchronization in alpha band coherence accompanied with enhanced gamma band coherence pinpoints alpha band coherence as signature of inter-individual variability. These observations further highlight a mechanistic difference in the processing of cross-modal stimuli between frequent and rare perceivers. Nonetheless, such differences are contingent on the stimulus as there was no difference in the global coherence pattern between frequent and rare perceivers during congruent /*ta*/. In retrospect of the global coherence patterns during McGurk stimuli, an obvious question is, do cross-frequency couplings among theta, alpha, beta and gamma band exist in a context specific way? Questions of such nature become a prime candidate to answer for future studies. A detailed account of cross-frequency coupling via coherence is currently out of scope of the present study.

### Characterization of NCN at source space

Pairwise coherence is affected by volume conduction to a considerable degree, specifically for local functional connectivity (Winter et al., 2007). The global coherence results are affected to a lesser degree by volume conduction, simply because the functional connections that can spuriously affect a distinct pattern of coherence are unlikely to survive the normalized vector summation procedure that is undertaken. Nonetheless, we need to validate if at least qualitatively the source and sensor level analysis are consistent. Subsequently, the global coherogram computed from reconstructed sources, first estimated through LCMV analysis were explored. The locations that showed statistical significance in the ratio of source power between /*ta*/ and /*pa*/ trials were used for reconstruction of sources. Frequent and rare perceivers showed a considerable overlap in brain areas involving right STS, fusiform gyrus, left inferior frontal gyrus and bilateral superior frontal gyrus. When coherogram was computed at the source level and the difference of global coherence between /*ta*/ and /*pa*/ are plotted, we could identify a high degree of similarity with the sensor space results (**Figure 4B-G**). Even though the exact spectro-temporal boundaries were slightly different, the overall pattern of results of enhanced gamma coherence and decreased alpha coherence at zero AV lag, and broadband coherence for ±450 ms AV lag was observed. Crucially, the major overlap of cortical sources across frequent and rare perceivers pinpoints the significance of understanding the communication within network of cortical regions over emphasizing role of isolated cortical loci in cognition

### Mechanistic understanding of NCN dynamics using biologically realistic computational model

Alpha and gamma band coherences are observed in processing of multisensory stimulus (Hummel & Gerloff, 2005; Kanayama et al., 2007; Doesburg et al. 2008; Kayser et al., 2008; Maier et al., 2008; Kayser & Logothetis, 2009; Kumar et al., 2016; also present results, **Figure 2**). Interestingly, high gamma coherence is seen when the nature of multisensory stimulus is complex (asynchronous, incongruent) (Doesburg et al., 2008; Kumar et al., 2016 and present results, **Figure 2**) which in some instances lead to illusory perception (Kanayama et al., 2007; Kumar et al., 2016; and present results, **Figure 1**). Gamma coherence is also observed in the communication involving higher order multisensory areas (Maier et al., 2008; and present results, **Figure 4**). Our computational model explains that alpha band coherence emerges when visual system has a direct influence on auditory node, while gamma coherence was observed only with indirect A-V interactions via multisensory node (**Figure 6 Supplement 1**). From a theoretical perspective this is possible because the time scale of processing is most disparate for the auditory and visual system, with auditory the fastest and visual the slowest. Without the presence of an intermediate time-scale, one “mode of communication” (alpha coherence) is sustained by the neural mass model within biologically relevant parameter regimes. Once there is another neural mass of intermediate time-scale participating in processing of information, the higher dimensionality of the resultant dynamical system allows creation of another mode of communication. Hence, our model suggests that gamma coherence could emerge due to the communication between primary auditory and visual areas but routed indirectly via higher order areas such as pSTS or inferor parietal or frontal areas. Our suggestion is in line with earlier observations of visual stimuli modulating auditory perception either directly resulting in alpha coherence (Kayser et al., 2008) or indirectly via higher order regions (STS) resulting primarily in gamma coherence (Maier et al., 2008; Kayser & Logothetis, 2009).

Behavioral responses from rare perceivers indicate limited influence of visual stimulus in shaping up the perceptual response since their response is akin to unisensory auditory response. The neuromarker of inter-individual variability, alpha coherence was drastically diminished (desynchronization) when A-V coupling was extremely weak (*W*_*AV*_ < 0.05) (**Figure 6A**). This indicates that overall interaction between visual and auditory node (direct and indirect via pSTS for example) is comparatively lesser in rare perceivers with respect to frequent perceivers and thus, results more in unisensory perception. Subsequently, we can also infer that direct A-V coupling is crucial for “frequently” perceiving the illusion of McGurk stimulus as in the case of frequent perceivers. On the other hand, differences in illusory perception and unisensory perception in both kinds of perceivers emerge from indirect A-V coupling via multisensory node (**Figure 6 B & C**). As discussed before, high gamma coherence is associated with multisensory processing involving interaction with higher order multisensory areas (Maier et al., 2008). Supporting this observation, we show A-V communication via multisensory node is crucial to generate gamma coherence during illusory perception in frequent and rare perceivers.

Alpha and/or gamma coherences have been observed in other Audio-Visual perception studies involving A-V speech phrases (Doesburg et al., 2008), natural A-V scenes (Kayser et al., 2008) and also artificially generated A-V looming signals (Maier et al., 2008). Increase in gamma coherence and reduction in alpha and beta coherences were observed during the perception of incongruent (lagged) A-V speech phrases (Doesburg et al., 2008). Increase in the interaction between fast and slow nodes via intermediate node increases the gamma coherence and decreases coherences in alpha and beta band (**Figure 6B**). Therefore, a similar mechanism that explains the observations of McGurk illusory perception is also applicable for explaining observations during perception of incongruent (lagged) A-V speech phrases. Increase in A-V interactions via multisensory node also explains the enhanced gamma coherence between auditory cortex and Superior Temporal Sulcus during congruent A-V looming signals in rhesus monkeys (Maier et al., 2008). Similarly, strong A-V interactions that distinguish the two kinds of perceiver groups (**Figure 6A**) also explain the increase in alpha phase consistency observed during natural A-V scenes in rhesus monkeys (Kayser et al., 2008). A different configuration of the model, where fast (auditory) node is source of EI and the slow (visual) node is sink of EI, generates peaks in beta band coherence (**Figure 6 Supplement 2B**) whereas the default configuration generates peaks in beta band as well as alpha band coherences (**Figure 6 Supplement 1B**). Therefore, this difference in configurations distinguishes visual perception of words (increase in beta band coherence and decrease in alpha band coherence) from auditory perception of words (increase in alpha and beta band coherence) suggesting that auditory (fast) node is the sink of EI during auditory perception and visual (slow) node is the sink of EI during visual perception (von Stein et al., 1999). Stretching to studies other than audio-visual perception, direct interactions between fast and slow nodes also explain the observed high alpha coherence during good performance while matching tactile Braille stimulus with its visual counterpart (Hummel & Gerloff, 2005) and the fast-slow indirect interactions via intermediate time-scale node explains the high gamma band coherence during rubber-hand illusion when visuo-tactile stimuli were congruent (Kanayama et al., 2007).

We have speculated the specific interactions of neural masses with different time-constants that generate band specific coherences and that are responsible for their enhancement and diminution. Multi-parametric and unbounded nature of the parameter space results in myriads of dynamics including chaos which is non-biological (Stefanescu & Jirsa, 2008). Therefore, such models should not be used to directly fit the data by estimating model parameters that minimize the error using optimization techniques. However, our model will be useful as a phenomological or minimalistic model in providing mechanistic insights into many findings (Fries, 2015; Engel et al., 2012) including pathological conditions (Başar & Güntekin, 2008) where relative changes in band specific coherences are observed.

## Materials and Methods

### Participants

Twenty nine normal healthy volunteers (16 males and 13 females, in the range of 21-29 years of age; mean age 25, SD = 3) participated in the study. All participants gave written informed consent in a format approved by the Institutional Human Ethics Committee of the National Brain Research Centre, Gurgaon which is in agreement with the Declaration of Helsinki. None of the participants had a history of neurological or audiological problems and were compensated for their time devoted to the experiment. All had normal or corrected-to-normal vision and were right-handed (tested using Edinburgh handedness inventory). The data from four volunteers were not included in the study because the channel impedance values in EEG exceeded 10 kΩ.

### Stimuli and trials

The experiment composed of 360 trials in which videos of a native Hindi speaking male articulating the syllables */ka/* and */ta/* (Fig. 1A) were presented. One-fourth (90 trials) of the trials consisted of congruent videos (visual */ta/* auditory */ta/*). The remaining three-fourths of the trials comprised incongruent videos (visual */ka/* auditory */pa/*) presented with AV lags: −450 ms (audio leads the articulation), 0 ms (synchronous) and +450 ms (articulation leads the audio), each encompassing one-fourth of the overall trials. The auditory object in the incongruent trials was extracted from a video of the speaker articulating */pa/* using the software Audacity (www.audacityteam.org). Subsequently, the extracted auditory /*pa*/ was superimposed onto the muted video of the speaker articulating the syllable */ka/* using the software Videopad Editor (www.nchsoftware.com). The composite multisensory stimuli were rendered into an 800 x 600 pixels movie with a digitization rate of 29.97 frames per second. Stereo soundtracks were digitized at 48 kHz with 32 bit resolution. Presentation software (Neurobehavioral System Inc.) was used to present the stimulus videos using a 17” LED monitor. Sound was delivered using sound tubes at an overall intensity of ∼60 dB.

### Experimental design

The experiment was divided into three blocks. Each block consisted of 120 trials comprising all the four kinds of videos (30 trials of each). Inter-stimulus intervals were pseudo-randomly varied between 1200 ms and 2800 ms to minimize expectancy effects. Using a forced choice task, the participants had to indicate their choice by pressing a specified key on the keyboard whether they heard */ta/, /pa/* or something else (others) while watching the videos.

### Eye Tracking

Gaze fixations of participants on the computer screen were recorded by EyeTribe eye tracking device (https://theeyetribe.com/). The gaze data were analyzed using customized MATLAB codes. The image frame of the speaker video was divided into 2 parts, the head, and the mouth. The gaze fixations at these locations over the duration of stimulus presentation were converted into percentage measures for further statistical analysis.

### EEG recording

Continuous EEG scans were acquired using a Neuroscan system (Synamps2, Compumedics, Inc.) with 64 Ag/AgCl scalp electrodes sintered on an elastic cap in a 10-20 montage. Recordings were made against the centre (near Cz) reference electrode on the Neuroscan cap and digitized at a sampling rate of 1000 Hz. Channel impedances were monitored to be at values < 5kΩ. Four volunteers showing higher impedances (∼10 kΩ) were discarded from further analysis.

### EEG Data processing

In the preprocessing step, the acquired EEG data was filtered using a band pass of 0.2-45 Hz. Subsequently, epochs of 900ms post the onset of first sensory object (auditory vocalization or articulatory lip movement) was extracted. Epochs extracted from congruent and incongruent videos were further sorted based on the perceptual experience: /*ta*/, /*pa*/ and ‘others’. The sorted epochs were then baseline corrected by removing the temporal mean of the EEG signal on an epoch-by-epoch basis. Finally, in order to remove the response contamination from ocular and muscle-related artifacts, epochs with maximum signal amplitude above 50 µV or a minimum below −50 µV were removed from all electrodes.

### Network analysis and global coherogram

To investigate frequency specific FC that subserves cross-modal perception and characterizes a frequent from a rare perceiver, we computed the global coherogram. Global coherogram captures the global coherence dynamics and quantifies the strength of neural co-activation across the whole brain at specific frequencies over time. In order to compute global coherogram from the preprocessed time series sorted based on the perceptual categories, we employed the Chronux (Mitra & Bokil, 2008) function cohgramc.m to obtain trial-wise time frequency cross-spectral matrix for all the sensor combinations. The output variable ‘S12’ of the function cohgramc.m yields the time frequency cross-spectrum density at a frequency *f* between sensor pair *i* and *j* employing the formula:

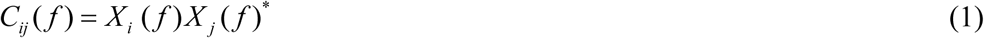

where, *C*_*ij*_ (*f*) represents the cross spectrum, *X*_*i*_ (*f*) represent the tapered Fourier transform of the time series from the sensor *i* and *X* _*j*_ (*f*)* represent the complex conjugate of the tapered time series from the sensor *j* at frequency *f*. In our analysis, a 62 x 62 matrix of cross spectra that represents all pairwise sensor combinations was computed. The time bandwidth product and the number of tapers were set at 3 and 5, respectively, and a moving window of 0.4 s with a step size of 0.05s were employed in the computation. Thereafter, we computed the global coherence at each time and frequency bin by computing the ratio of the largest eigenvalue of the cross-spectral matrix to the sum of the eigenvalues on a trial-by-trial basis employing the following equation:

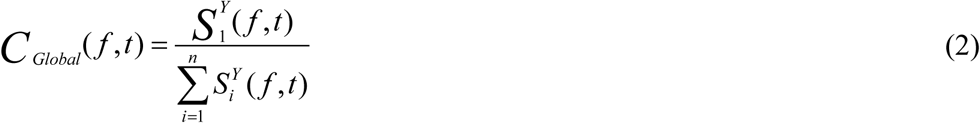

Where *CGlobal* (*f*, *t*) represent the global coherence at frequency *f* in the time window *t*, 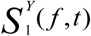 represent the largest eigenvalue and the denominator 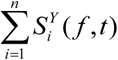 represents the sum of eigenvalues of the cross-spectral matrix at every time bin. Subsequently, the time-frequency global coherogram computed for */ta/* and */pa/* responses were compared non-parametrically using cluster based permutation tests for frequent and rare perceivers explicitly (Maris et. al., 2007; Kumar et. al., 2016).

We computed the global coherence collapsed across the entire epoch to identify if there are certain frequencies around which the network is most robust underlying cross-modal (illusory /*ta*/) and unimodal (/*pa*/) perception in frequent and rare perceivers. Furthermore, to investigate whether the organization of these networks dependent on the stimulus configurations or the perceptual outcome, we also computed the global coherence during congruent /*ta*/ perception in frequent and rare perceivers. We employed the Chronux function CrossSpecMatc.m for computing the global coherence. The output variable ‘Ctot’ of the function yields the global coherence value at frequency *f* by initially computing the cross-spectrum for all sensor combinations following the Equation 1. Subsequently, global coherence at every frequency bin is obtained by computing the ratio of the largest eigenvalue of the cross-spectral matrix to the sum of the eigenvalues on a trial-by-trial basis employing Equation 2. The time bandwidth product and the number of tapers were set at 3 and 5, respectively, and a fixed window size of 0.9 s was employed in the computation. Finally, we employed Silvermann’s bootstrapping test for detecting the presence of a bimodal distribution (Silverman, 1981). We performed Silvermann’s bootstrapping bimodality test on the time averaged global coherence separately on the perceptual categories across all AV lags in frequent and rare perceivers.

The aforementioned analysis was further performed to compute the global coherogram and coherence during the perception of congruent /*ta*/ in frequent and rare perceivers. Subsequently, the global coherogram was compared employing cluster based permutation tests.

### Source Reconstruction and functional connectivity

To investigate if the global coherogram patterns observed at the sensor level affected by volume conduction, we constructed source time-series and computed the global coherogram differences between /*ta*/ and /*pa*/ at all AV lags in frequent and rare perceivers. We employed a linearly constrained minimum variance (LCMV) beamformer algorithm (Van Veen et al., 1997) to identify the cortical generators of the time-series during /*ta*/ and /*pa*/ perception in frequent and rare perceivers. The entire epoch of 0.9s was employed in the source analysis. Prior to source reconstruction, we constructed our personalized average template from the individual MRIs of the subjects using the function ‘antsMultivariateTemplateConstruction’ developed by Advanced Normalization Tools (ANTs) (http://stnava.github.io/ANTs/)(Avants et al., 2011). The pipeline initially involves rigidly registering the participants T1 images to a MNI template while maintaining the volume and size of the original structural images. The rigidly registered images are then averaged to generate a temporary template. This template is then used as the first registration target onto which each participants T1 image is non-linearly registered, transformed and averaged. Iteratively, the T1 images are non-linearly registered to the new average, transformed and re-averaged generating a relatively a more precise average for the next iteration.

For source reconstruction we employed Fieldtrip toolbox. Firstly, we used ft_prepare_leadfeild.m and employed the Boundary Element Method (BEM) to generate the leadfield matrix from the template we constructed. The leadfield matrix corresponds to the tissue and geometrical properties of the brain represented as discrete grids or voxels. Subsequently, we employed ft_timelockanalysis.m to evaluate the covariance matrix of the epochs sorted based on perceptual categories in frequent and rare perceivers as the LCMV adaptive spatial filters are constrained by the covariance and leadfield matrices. These spatial filters regulate the amplitude of brain electrical activity passing from a specific location while attenuating activity originating at other locations. The distribution of the output amplitude of the spatial filters provides the metric for source localization. However, in order to compare the source power during /*ta*/ and /*pa*/ perception, we computed an inverse "common spatial filter? employing ft_sourceanalysis.m from the dataset obtained by appending the datasets of /*ta*/ and /*pa*/ post time-lock analysis. Eventually, based on the pre-computed common spatial filter we evaluated the sources separately for /*ta*/ and /*pa*/ employing ft_sourceanalysis.m. The difference in the source power between /*ta*/ and /*pa*/ were consequently compared by taking the ratio of the source power of /*ta*/ and /*pa*/. Finally, the grids eliciting power above the 99.5th percentile were identified as sources and were interpolated onto the constructed template for illustrative purposes.

For reconstructing time series from the thresholded sources, we projected single trial epoched time series from sensors onto the source space by multiplying them to the concordant spatial filters of the thresholded sources. There were overall 52 grids of the spatial filter corresponding to the sources represented in **Figure 4A** onto which the sensor level data was projected to obtain the source time series. Furthermore, each spatial filter is represented by three components representing the unity moment in the *x, y* and *z* direction of the dipole at the respective grid location. We estimated the global coherogram differences between the /*ta*/ and /*pa*/ perception in frequent and rare perceivers from the source time series from the component that best matched the sensor level global coherogram results.

### Large scale dynamical model of three neural masses

Our objective was to construct a large-scale dynamical model which is biologically realistic to explain the generative mechanisms underlying observed coherence spectra and frequency specific functional connectivity during illusory and non-illusory perception in rare and frequent perceivers based on empirical data. Our proposed model is a network of three neural masses, each comprising of excitatory and inhibitory neurons representing auditory, visual and higher order multisensory cortical regions (**Figure 5**). We follow a previously established practice and convention in computational modelling by treating each cortical region as an individual node as suggested by Stefenascu and Jirsa (Stefanescu & Jirsa, 2008).

Broadly we incorporate the following biophysically realistic factors in our model construction.

1. The time-scale of processing of the visual system can be considered slowly varying in comparison to auditory system (Williams et al., 2004; Rosen & Howell, 2011). Multisensory system can be placed in between the auditory and visual systems in terms of the processing time-scale.
2. Two of the ways visual inputs are directed to the auditory cortex are: 1) visual cortex could directly influence the auditory cortex in a feedforward manner due to direct projections (Falchier, et al., 2002; Rockland & Ojima, 2003; Wallace et al., 2004) and 2) feedback from the higher multisensory association areas (Bizley & King, 2012). Hence, in our proposed model visual node influences the auditory node in both manners: directly and indirectly via multisensory node.
3. As post-synaptic potentials of pyramidal cells, which are excitatory, are shaped by their connections with other excitatory cells and inhibitory cells (Kirschstein & Köhling, 2009). We use a population of excitatory and inhibitory neurons in each node where the number of excitatory neurons are considerably higher (Olbrich & Braak, 1985). Thus, 150 excitatory neurons and 50 inhibitory neurons are selected to have a 3:1 ratio between them, an approach previously followed by Stefanescu and Jirsa (Stefanescu & Jirsa, 2008). Inhibitory neurons in one neural area do not directly influence inhibitory neurons within the same area since such connections are sparse in nature (Wilson & Cowan, 1972; Stefanescu & Jirsa, 2008).

Incorporating these factors we define a dynamic mean field model that comprises of three equations for an excitatory Hindmarsh Rose (HR) neuron (number of excitatory neurons are 150 within an area, *N*_*E*_ = 150) and three equations for an inhibitory HR neuron (number of inhibitory neurons are 50 within an area, *N*_*I*_ = 50) (**Figure 5**). The three variables account for the membrane dynamics and two kinds of gating currents, one fast and one slow respectively. Thus, the entire network can be represented as a network of coupled non-linear differential equations comprising of

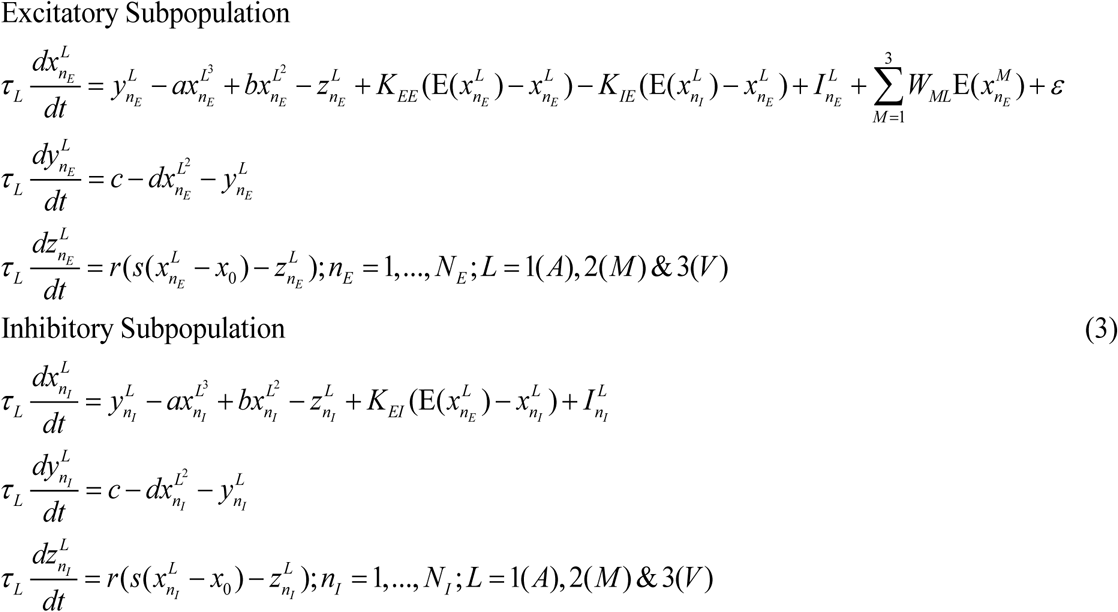

Where *L*: A, V and AV for auditory, visual and audio-visual areas that are driven by a common noise distribution (*ε*). In our model auditory node has the fastest time-constant (*τ* _*A*_ ∼ 0.05*ms*), visual node has the slowest time-constant (*τ*_*V*_ ∼ 2.5*ms*) and time-constant of multisensory node is chosen to be in between the two (*τ*_*M*_ ∼ 1*ms*) as it integrates information from both the modalities. The mean activity of excitatory neurons in a node (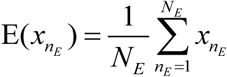 influences neuronal activities of other nodes that is governed by coupling parameters: *W*_*AV*_ (auditory-visual coupling), *W*_*AM*_ (auditory-multisensory coupling) and *W*_*VM*_ (visual-multisensory coupling). Positive value of coupling parameters reflects excitatory influence and negative value reflects inhibitory influence. Inhibitory influences are chosen to maintain a balance with excitation. For example, visual node’s excitatory influence of +*W*_*AV*_ on auditory node is balanced with inhibitory influence of the same strength (-*W*_*AV*_) from the auditory node.

In this configuration, visual node is referred as source node as it is the source of excitatory influence whereas auditory node is referred to as sink node as all excitatory influences are directed towards auditory node and multisensory node behaves as both source and sink..

We place each individual neuron in a dynamical regime where both spiking and bursting behavior is possible depending on the external input current (I) that enters the neuron when other parameters are held constant at the following values: *a* =1;*b* = 3;*c* =1; *d* = 5; *s* = 4; *r* = 0.006; *x*_0_ = −1.6; (Stefanescu & Jirsa, 2008).

The coupling between the neurons within a node is linear and its strength is governed by the following parameters: *K*_*EE*_ for excitatory-excitatory coupling,*K*_*EI*_ for excitatory-inhibitory coupling and *K*_*IE*_ for inhibitory-excitatory coupling. As excitatory and inhibitory synapses are not independent processes, their relation is captured by the ratio 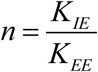.As alpha (8-12 Hz) and delta (1-4 Hz) rhythms are observed during resting state (Gold et al., 2006), the inhibition to excitation ratio (*n* = 3.39) is chosen when the average activity of nodes in a disconnected network has higher power at alpha and delta frequencies in the absence of stimulus (*μ*(*I*_*A,V*, *M*_) = 0.1; baseline) (**Figure 5 Supplement 1**). The external currents to both the excitatory and inhibitory subpopulations are drawn from a Gaussian distribution where *μ* and *σ* are the mean and standard-deviation. As the input stimulus relays to auditory, visual and multisensory regions via thalamus, we interpret lateral geniculate nucleus (LGN) and medial geniculate nucleus (MGN) to be the source of external current (*I*_*A*_, *I*_*V*_ and *I*_*M*_) pulse of 450 ms in the nodes when the model was simulated for 1 sec. In rhesus monkey, the projections of MGN to pSTS were found to be sparse (Yeterian & Pandya, 1989). Therefore, we choose lower mean value of external current to multisensory node (*μ*(*I*_*M*_) = 0.85) in comparison to visual node (*μ*(*I*_*V*_) = 2.8) and auditory node (*μ*(*I*_*A*_) = 2.8) while keeping the standard deviation of the external current at 0.4 for all nodes.

## Acknowledgements

The study was supported by NBRC Core funds and by grants Ramalingaswami fellowship, (BT/RLF/Re-entry/31/2011) and Innovative Young Bio-technologist Award (IYBA), (BT/07/IYBA/2013) from the Department of Biotechnology (DBT), Ministry of Science and Technology, Government of India to AB. AB also acknowledges the support of Centre of Excellence in Epilepsy and MEG (BT/01/COE/09/08/2011) from DBT. DR was supported by the Ramalingaswami fellowship (BT/RLF/Re-entry/07/2014) from DBT.

---

**Figure 1 Supplement 1:**
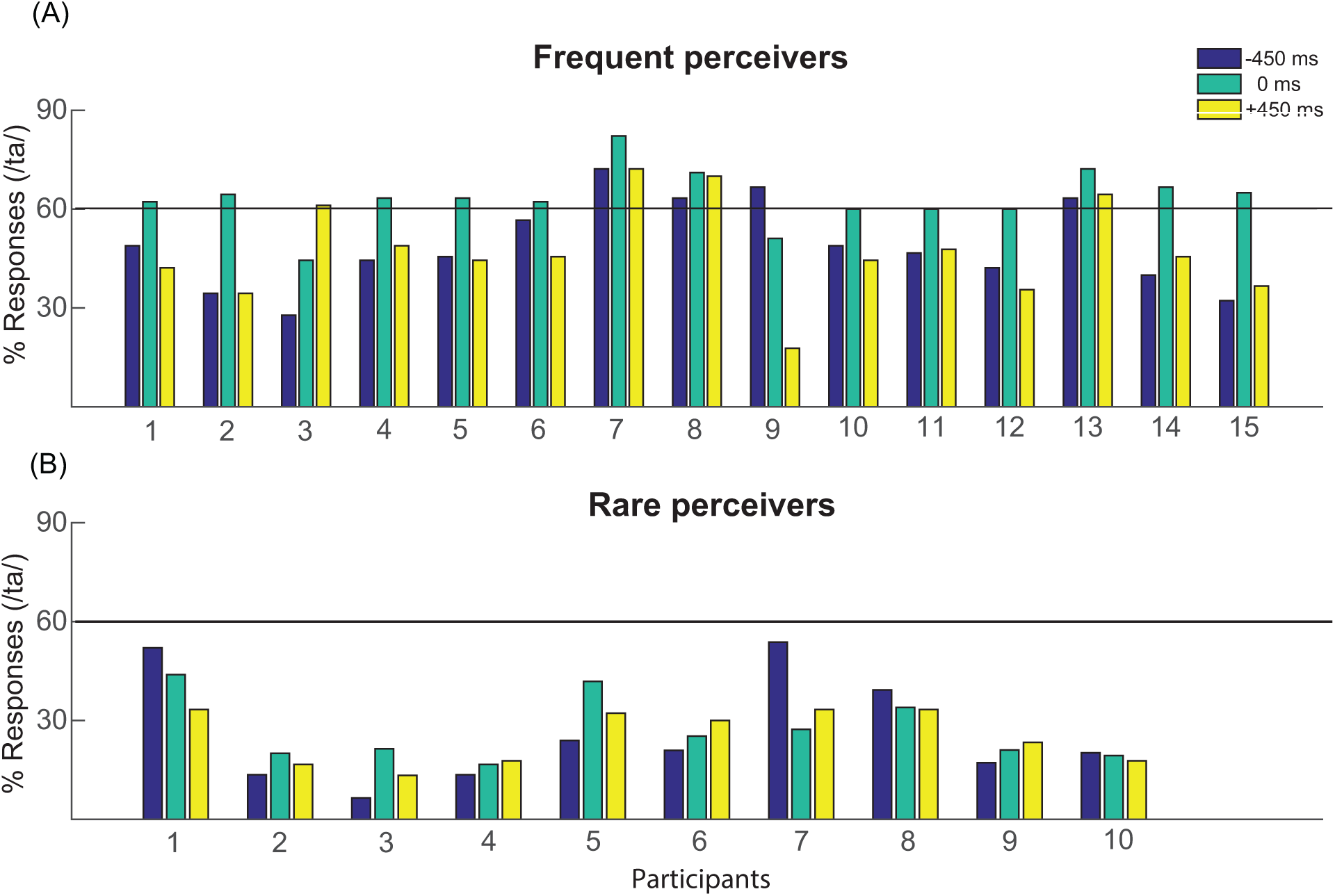
Individualist participant behavior: Percentage of /ta/ responses during the 0 ms and 450 ms AV stimuli in (A) Frequent perceivers (B) Rare perceivers.

**Figure 1 Supplement 2:**
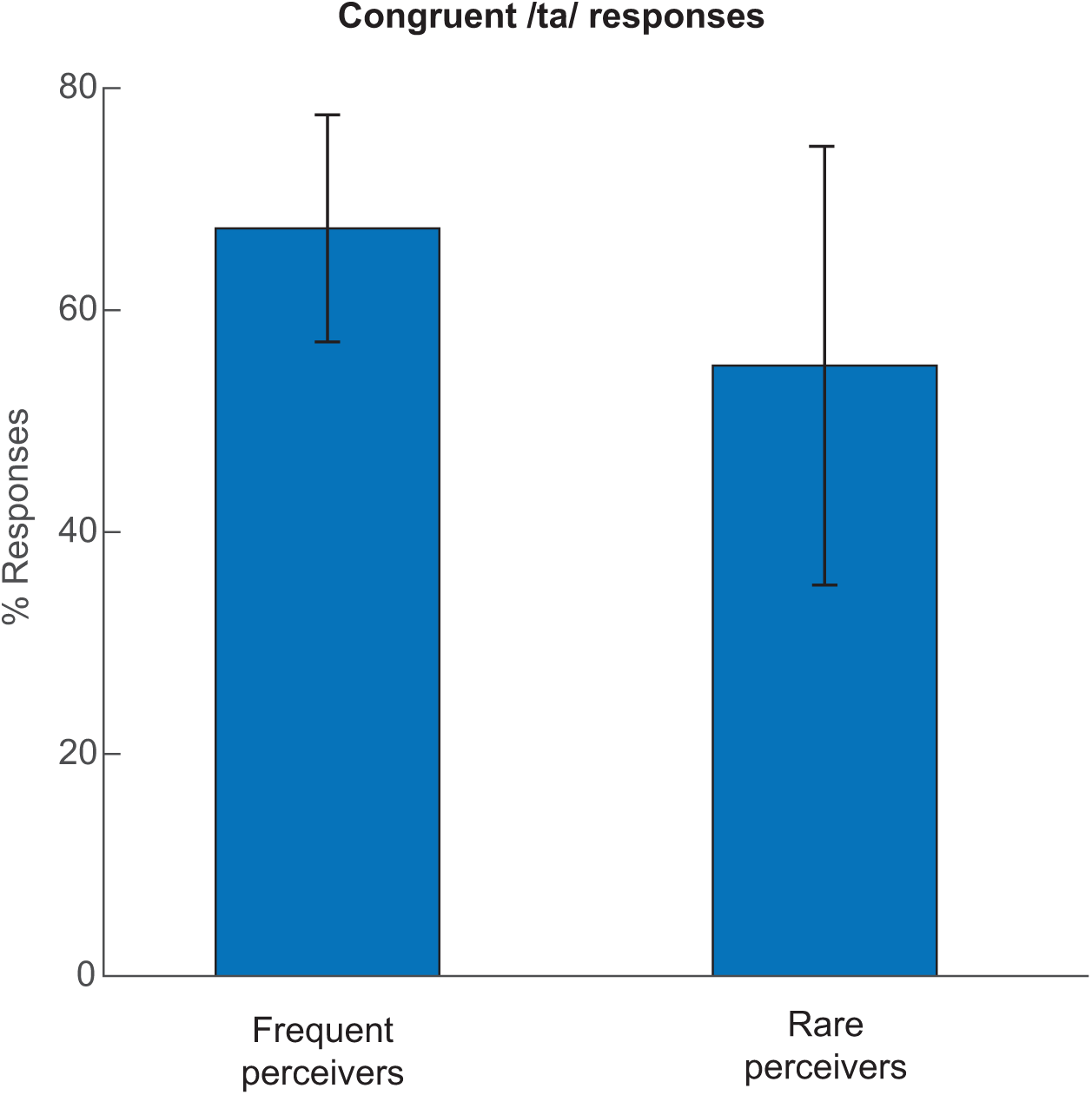
Hit rate during congruent /ta/ stimulus: The percentage of /ts/ responses trail-by-trial across the participants.

**Figure 1 Supplement 3:**
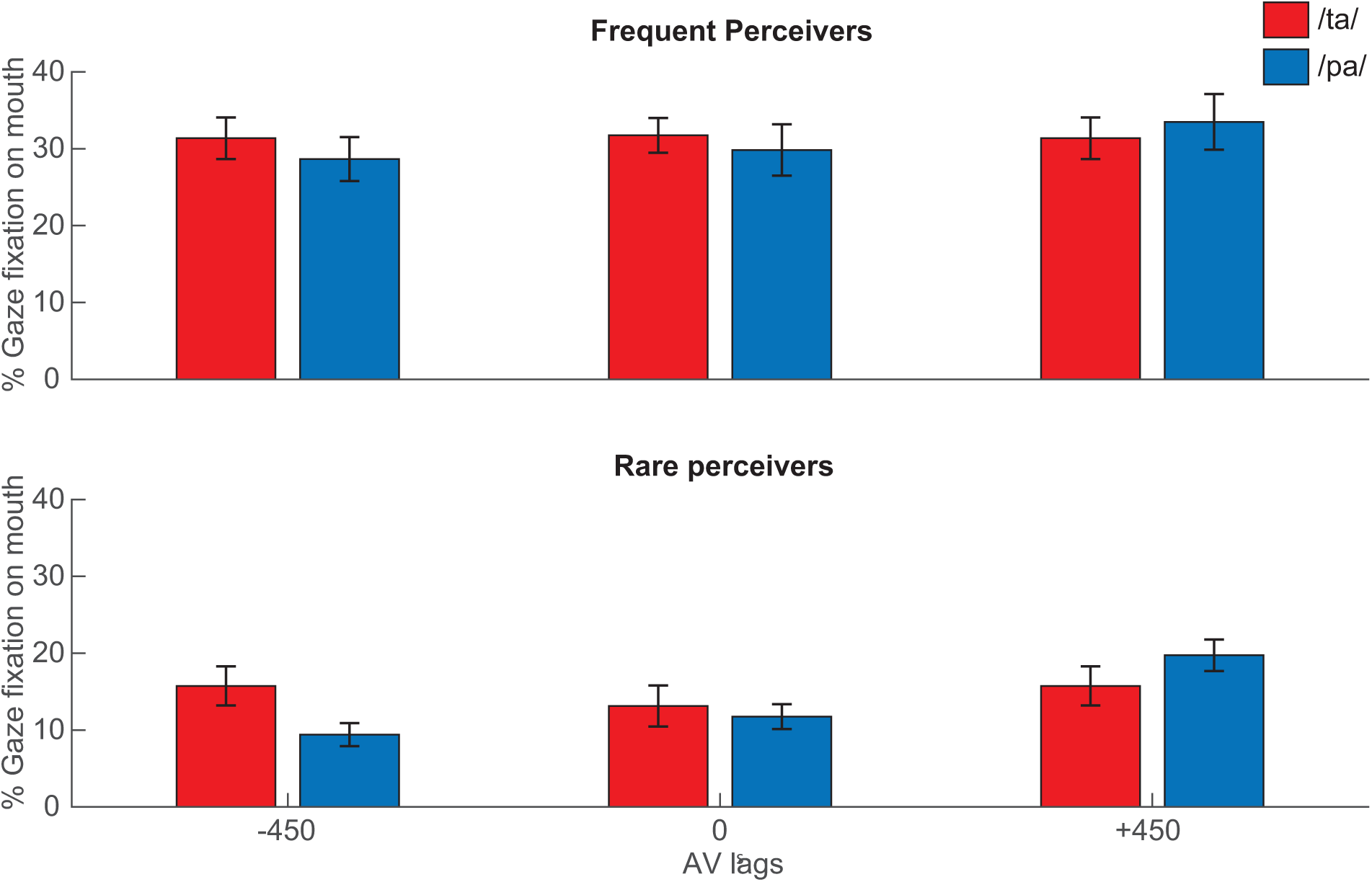
Gaze behavior: Percentage of gaze fixations on the mouth of the articulator in the AV stimuli averaged trail-by-trial across the participants (A) Frequent perceivers (B) Rare perceivers.

**Figure 5 Supplement 1:**
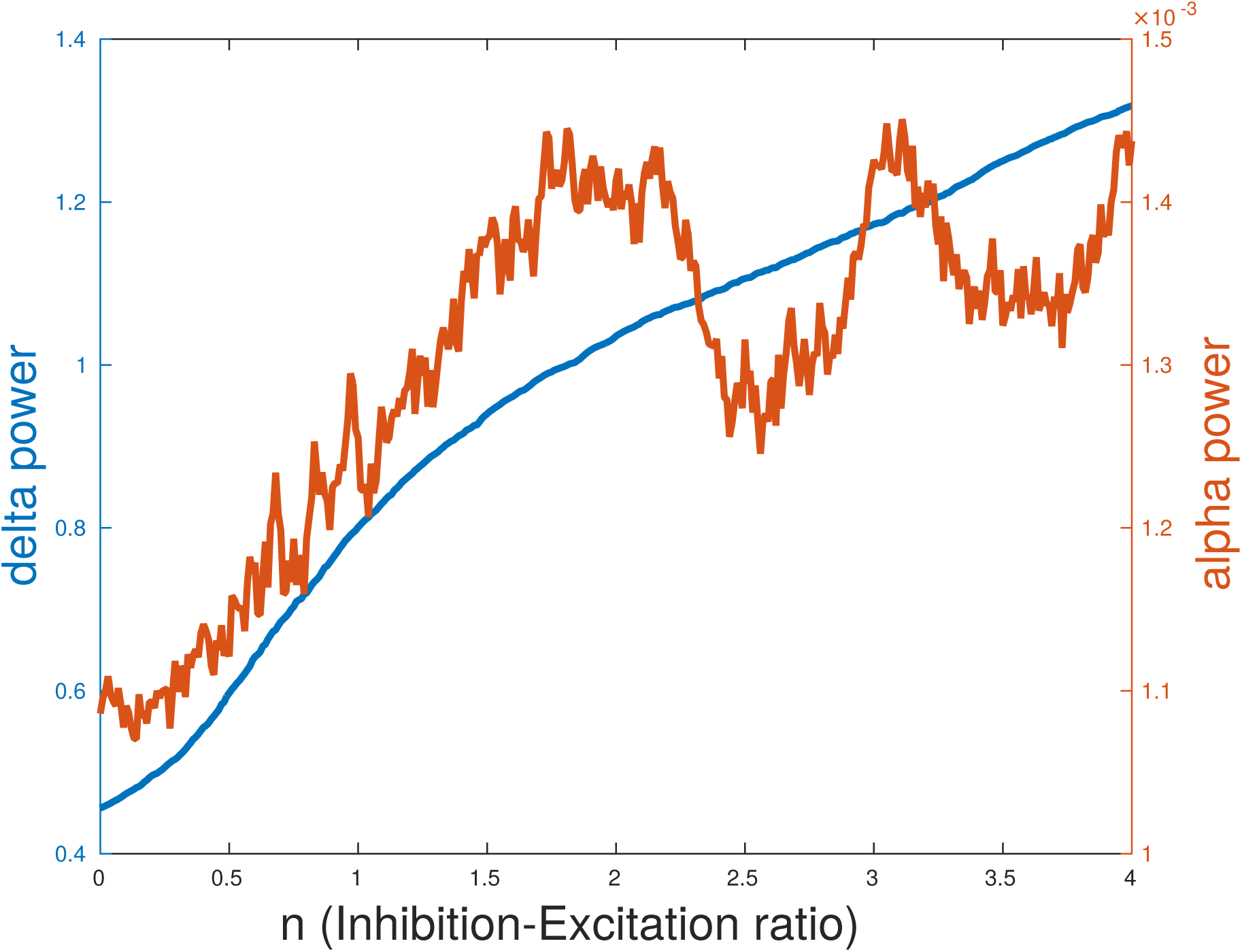
Selection of Inhibition-Excitation ratio (n): Delta and alpha power when the nodes are disconnected and driven by baseline current (I=0.1). Selection of n was made in order to have comparatively higher delta and alpha power.

**Figure 6 Supplement 1:**
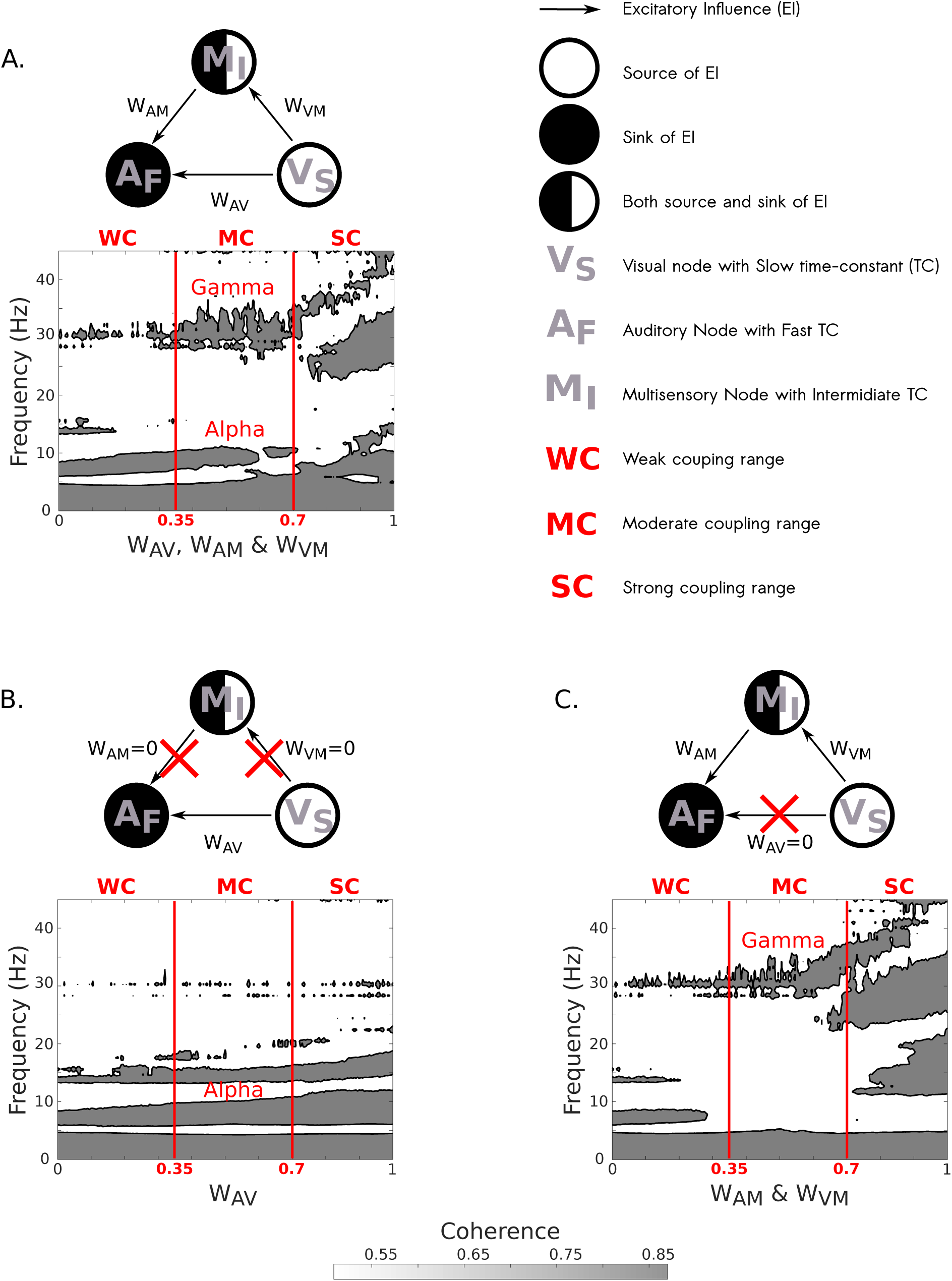
Prediction of alpha and gamma coherences from neural mass model: A) Alpha and gamma band coherence co-exist in moderate coupling range. B) Only direct A-V coupling generates alpha coherence independently. C) Indirect A-V coupling via multisensory node generates gamma coherence at the limit case scenario of weak direct coupling.

**Figure 6 Supplement 2:**
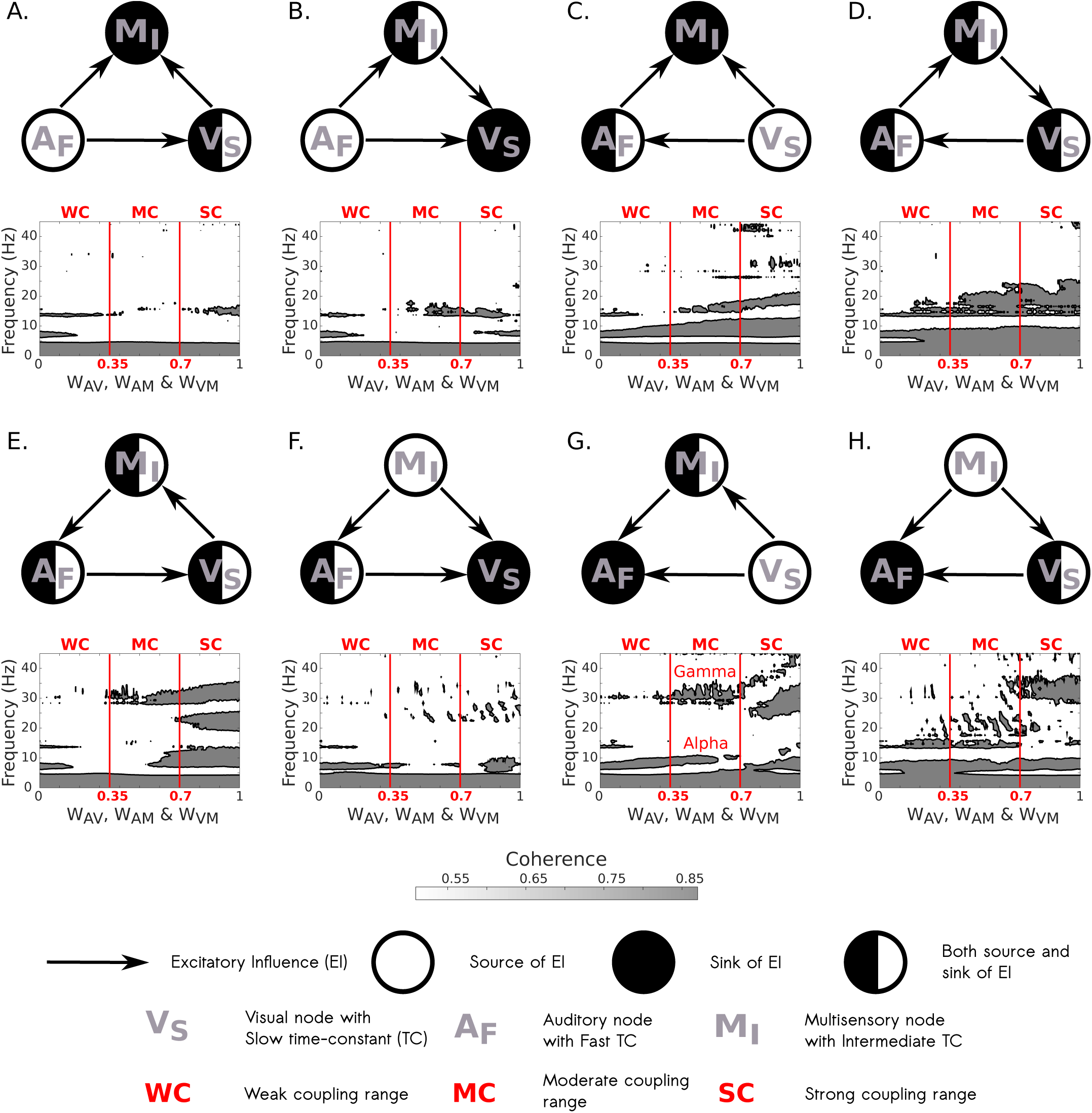
Global coherence for different source-sink combinations: G) Only when visual node is source and auditory node is the sink (as in our model), we observe co-existence of alpha and gamma band coherence in moderate coupling range. A)-F) and H) Exploration of various coupling scenarios to identify if it is possible to generate alpha and gamma coherence in moderate coupling range.

